# *Ficd* loss rescues motor impairments and reverses oligodendrocyte maturation deficits in a mouse model of spinocerebellar ataxia type 3

**DOI:** 10.64898/2026.08.07.743629

**Authors:** Kate M. Van Pelt, Yamei Deng, Alexey I. Nesvizhskii, Henry L. Paulson, Maria do Carmo Costa, Matthias C. Truttmann

## Abstract

Spinocerebellar ataxia type 3 (SCA3) is an inherited, fatal neurodegenerative disease caused by a pathological CAG repeat expansion in the ATXN3 gene, resulting in the selective degeneration of vulnerable neuronal populations. Recent work has identified impairments in oligodendrocyte maturation as a novel and robust feature of SCA3 pathogenesis. Oligodendrocytes synthesize myelin structural components through the endoplasmic reticulum (ER), rendering this organelle essential for white matter integrity. Despite this, the role of ER function in SCA3 remains unclear. In this study, we show that loss of FICD-mediated AMPylation, a post-translational modification regulating the ER-resident HSP70 chaperone, BiP, rescues motor impairments in a transgenic SCA3 mouse model. *Ficd*^-/-^ SCA3 mice exhibit significantly reduced levels of nuclear ATXN3 in vulnerable brain regions, while *Ficd*^+/+^ littermates show an increased burden of AMPylated BiP in the spinal cord, identifying aberrant AMPylation as a novel contributor of SCA3 pathology. Using unbiased proteomics, we demonstrate that *Ficd* deletion mitigates the pathological decrease in myelin structural proteins and oligodendrocyte maturation factors, restoring levels of mature, myelinating oligodendrocytes. In parallel, we show that *Ficd* activates SREBP2-dependent cholesterol biosynthesis to support myelination. Taken as a whole, these findings posit ER homeostasis as a critical driver of oligodendrocyte pathology and identify FICD as a novel target for alleviating non-neuronal toxicity in SCA3.

## INTRODUCTION

Spinocerebellar ataxia type 3 (SCA3), also known as Machado-Joseph disease (MJD), is the most common dominantly-inherited ataxia and the second most common polyglutamine (polyQ) expansion disease behind Huntington’s disease (*1*, *2*). SCA3 is caused by a pathological CAG repeat expansion in the *ATXN3* gene, which results in the production of a mutant ATXN3 protein containing an expanded polyQ tract (*3*). Patients affected by the disease exhibit progressive ataxia accompanied by dystonia, distal muscle atrophy, vestibular and speech difficulties, and oculomotor dysfunction (*4*). This observed decline in motor control is owed to selective neurodegeneration of, predominantly, the cerebellum, brainstem, basal ganglia, and the spinal cord (*5*, *6*). As a neurodegenerative disease, SCA3 is ultimately fatal, with death occurring 10-20 years after symptom onset (*7*). Despite significant progress in recent years towards understanding the cellular pathways dysregulated in SCA3 (*8*), there are currently no disease-modifying therapies available for this fatal disease.

ATXN3 is ubiquitously expressed in all cell types and localizes to both the cytoplasm and nucleus, where it functions as a deubiquitinase (*9*, *10*). In SCA3, mutant ATXN3 is mislocalized and forms aggregates in the nucleus, a pattern observed across polyQ diseases (*11*, *12*). Recent studies profiling transcriptomic changes in vulnerable brain regions from SCA3 transgenic and knock-in mouse models revealed significant alterations in gene expression in the presence of mutant ATXN3 (*13–16*). In contrast, loss of ATXN3 has minimal impact on gene expression (*14*) and *ATXN3* null mice present phenotypically normal (*17*). Together, these findings support the view that pathologically-expanded ATXN3 incites gain-of-function toxicity in SCA3.

As selective neuronal loss is the defining feature of SCA3 and related diseases, research in the field has historically taken a neuron-centric approach. Interestingly, however, white matter pathology has recently emerged as a prominent early and progressive feature of SCA3 pathogenesis across multiple studies in SCA3 mouse models (*13–16*, *18*, *19*). SCA3 transgenic mice exhibit pronounced downregulation of oligodendrocyte maturation genes in vulnerable brain regions, correlating with thinner myelination, reduced protein levels of myelin structural components, and a decrease in the number of mature, myelinating oligodendrocytes (*15*, *18*). The clinical relevance of these findings is corroborated by numerous imaging studies demonstrating early changes and atrophy in the cerebellar white matter tracts of SCA3 patients (*20–24*), as well as reduced myelin levels in postmortem tissue from SCA3 patients (*18*, *25*). Preclinical work has shown that treatment with anti-*ATXN3* antisense oligonucleotides (ASOs) is sufficient to provide functional rescue of this maturation deficit, but also resulted in undesirable hypermyelination in less-affected brain regions (*26*). Given that the mechanisms driving white matter pathology in SCA3 are unknown, additional research is needed to identify and characterize druggable candidates for the development of targeted therapies.

Myelin is a lipid-rich multilamellar sheath that insulates axons and is essential for both saltatory conduction and maintaining axonal integrity. With a single oligodendrocyte (OL) capable of myelinating up to 50 axons (*27*), these cells meet this high demand by synthesizing large amounts of lipids and proteins in the endoplasmic reticulum (ER) (*28*). As such, OLs are highly sensitive to disruptions to the secretory pathway and, in particular, ER homeostasis (*29*), rendering the ER a compelling focus for investigation into the cause of OL deficits in SCA3. Protein AMPylation, a post-translational modification (PTM) characterized by the addition of an adenosine monophosphate (AMP) group to serine or threonine side chains on target proteins, has recently emerged as a novel regulator of ER homeostasis (*30–35*). This PTM is performed by the fic domain-containing enzyme, FICD, which acts bi-functionally to catalyze both the addition (AMPylation) and removal (deAMPylation) of AMP to substrates (*36–38*). The most well-studied AMPylation target is the ER-resident HSP70 family chaperone, BiP, which is AMPylated in its substrate-free conformation, preventing ATP hydrolysis and subsequent client binding (*32*, *39*). Here, AMPylation is thought to “lock” BiP in an activated, ATP-bound state such that it can immediately engage with client proteins upon deAMPylation (*38*). In human patients, mutations in the active site of FICD have been linked to infancy-onset diabetes (Arg371Ser) and motor neuron disease (Arg374His) stemming from impaired deAMPylation activity, resulting in abnormally increased levels of AMPylated, inactive BiP (*40–43*). In contrast, loss of FICD is well-tolerated in the absence of stress in several model organisms, and is protective in stressor-specific contexts (*30*, *35*, *44–46*). We recently reported that *Ficd* knock-out mice are protected from pressure overload-induced heart failure through activation of the unfolded protein response in the ER (UPR^ER^) and ER-phagy (*47*). Changes in AMPylation levels have also been shown to modulate aggregation of several neurodegenerative disease-associated proteins (*48*), including α-synuclein (*49*, *50*), and loss of the *C. elegans* FICD ortholog, *fic-1*, suppresses polyQ toxicity via the UPR^ER^ (*51*). Still, whether attenuating FICD activity might be beneficial in a more physiologically-relevant polyglutamine disease model has yet to be explored.

Given the recent focus on oligodendrocyte pathology in SCA3 and literature identifying BiP/Grp78 as essential for the survival of myelinating cells (*52*), we asked whether the loss of FICD-mediated AMPylation attenuates behavioral and molecular signatures of mutant ATXN3 pathology. Using YAC-Q84 transgenic SCA3 mice, we show that whole-body *Ficd* deletion leads to a dramatic rescue of motor pathology in homozygous Q84/Q84 mice assessed over a year of motor and behavioral testing. At the molecular level, we correlate these findings with evidence that *Ficd*^-/-^ mice show significant reductions in ATXN3 nuclear accumulation, and identify a pathological increase in AMPylation in the spinal cord of Q84/Q84 mice as a potential novel driver of SCA3 pathology. Using data-independent acquisition mass spectrometry (DIA-MS), we analyzed the brainstem proteome of controls and Q84/Q84; *Ficd*^+/+^ and *Ficd*^-/-^ mice to reveal that *Ficd* deletion reshapes the myelination landscape - rescuing levels of myelin structural proteins, oligodendrocyte maturation factors, and key proteins involved in lipid synthesis. Corroborating previous reports, we show that Q84/Q84 mice exhibit significantly lower levels of mature, myelinating oligodendrocytes (CC1^+^/OLIG2^+^ double-positive cells) compared to WT/WT littermates, and that these cell populations are restored in *Ficd* null animals. Lastly, we use bulk RNAseq to identify increased SREBP2-dependent cholesterol synthesis in *Ficd*^-/-^ mice as a parallel pathway supporting oligodendrocyte health and myelin production. Taken collectively, these findings demonstrate that *Ficd* deletion rescues human ATXN3 pathology in a preclinical animal model of SCA3 and identifies myelination and lipid metabolism as key areas for exploring the role of small molecule *Ficd* inhibitors in therapeutic development for SCA3.

## RESULTS

### *Ficd* deletion rescues motor phenotypes in the YAC-Q84 mouse model of SCA3

In prior studies, we reported that the loss of FICD-mediated AMPylation in *C. elegans* alters polyglutamine (polyQ) aggregation dynamics (*48*) and suppresses their toxicity through the induction of a UPR^ER^-dependent protective signaling axis (*51*). While intriguing, the translatability of these results is limited by the expression of a pure polyQ repeat devoid of protein context in a non-neuronal cell type (*53*, *54*). We thus asked whether the loss of AMPylation could rescue polyQ disease phenotypes in the YAC-Q84 transgenic mouse model of spinocerebellar ataxia type 3 (SCA3). These animals express the full-length human *ATXN3* gene (*hATXN3*) harboring a pathogenic (CAG)_84_ repeat at near-endogenous levels and exhibit progressive motor impairment accompanied by mutant *ATXN3* neuropathological features (*55*). In contrast to other murine models of neurodegenerative disease, which can take months to develop disease-related behavioral changes, homozygous YAC-Q84 mice exhibit motor impairments as early as 4-6 weeks of age, rendering them a suitable choice for investigating the impact of *Ficd* depletion (*56*). As constitutive *Ficd* knock-out mice are viable with no overt phenotypes (*44*), we crossed *Ficd*^-/-^ and YAC hemizygous (Q84/WT) mice to obtain Q84/WT; *Ficd*^+/-^ breeders and established a colony. All experiments were conducted using YAC homozygous (Q84/Q84) *Ficd*^+/+^ or *Ficd*^-/-^ animals and corresponding YAC-negative (WT/WT) littermate controls (Fig. 1A, Fig. S1A-B). Across all time points, Q84/Q84 mice exhibited lower overall body mass relative to WT/WT controls, as reported previously (*56*, *57*), and we observed no effect of *Ficd* deletion on body weight regardless of sex or transgene status (Fig. S2A-B). Similarly, we found that *Ficd* genotype had no significant impact on CAG repeat size measured in the mouse tail (Supplementary Tables S2-S3), or brainstem transcript levels of endogenous mouse *Atxn3* (Fig. S1E-F) and the *hATXN3* transgene (Fig. S1G-H). Q84/Q84; *Ficd*^+/+^ animals exhibited comparable *Ficd* brainstem expression levels relative to WT/WT; *Ficd*^+/+^ littermates throughout the duration of the study (Fig. S1C-D), indicating that *Ficd* expression is unaffected in the presence of mutant hATXN3.

**Fig. 1.**
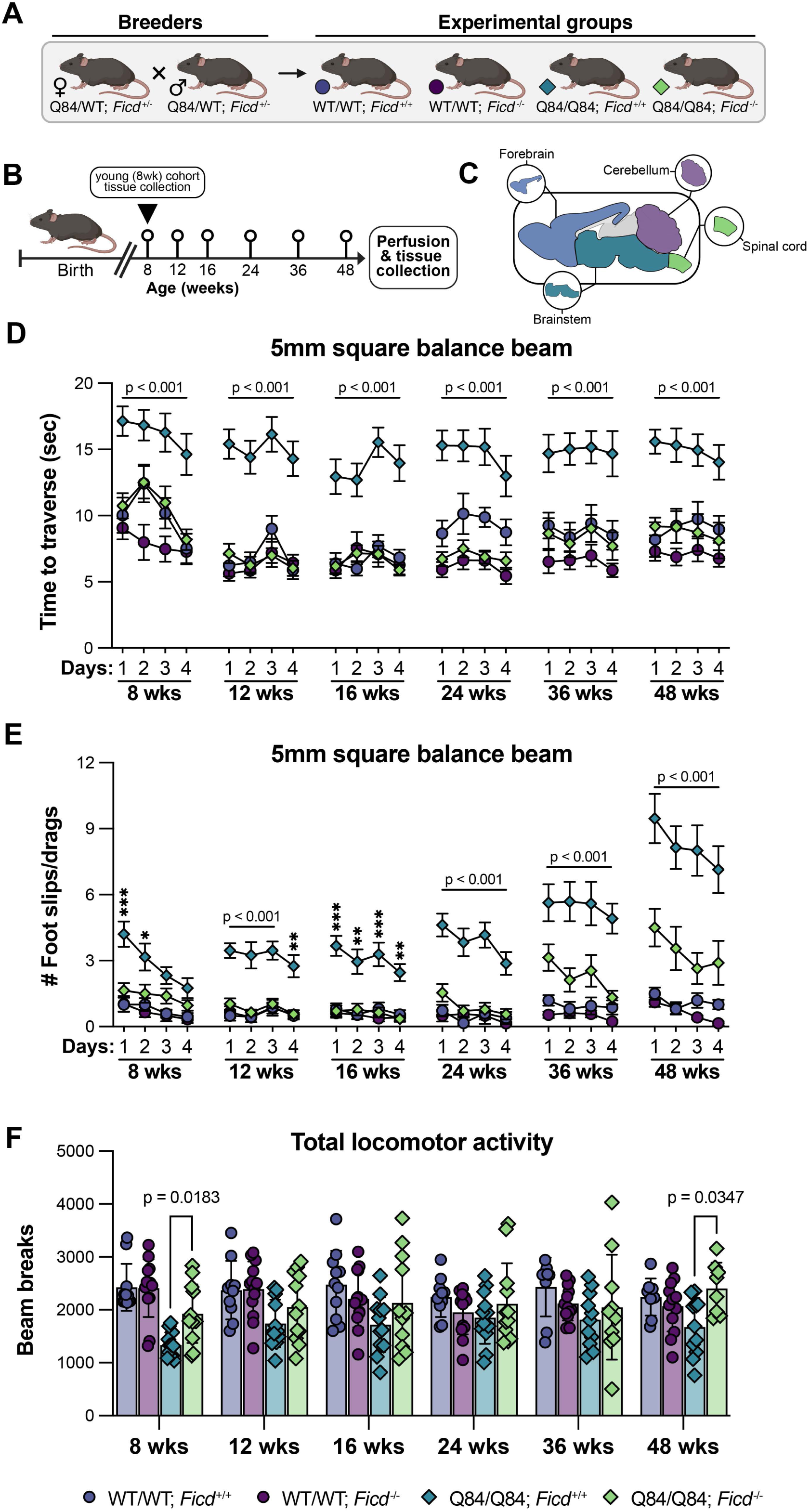
*Ficd* deletion rescues motor phenotypes in the YAC-Q84 mouse model of SCA3. (A) Breeding scheme employed in this study. YAC-hemizygous mice heterozygous for the *Ficd* null allele (Q84/WT; *Ficd*^+/-^) were bred to obtain YAC-negative (WT/WT) and YAC-homozygous (Q84/Q84) *Ficd*^+/+^ and *Ficd*^-/-^ experimental groups. (B) Experimental timeline. Mice were subjected to balance beam and open field assessments beginning at 8 weeks of age, and subsequently at 12, 16, 24, 36, and 48 weeks of age. Following conclusion of behavioral testing, animals were perfused and tissues were harvested for analysis. A separate cohort of 8 week-old mice was collected in the same manner. (C) Schematic depicting the brain regions of interest isolated by macro-dissection for biochemical analysis. (D) Average time taken to traverse the 5mm square balance beam on each of 4 consecutive trial days for all time-points tested. (E) Average number of hindlimb foot slips and/or drags during the 5mm square balance beam traversal task. For (D-E), data are displayed as mean ± SEM. (F) Total locomotor activity determined by the number of beam breaks recorded during open field testing (30 min), displayed as mean ± SD. For (D-E), statistical significance was determined by two-way ANOVA followed by Tukey’s post-hoc multiple comparisons testing. P-values reflect comparisons between Q84/Q84; *Ficd*^+/+^ and Q84/Q84; *Ficd*^-/-^ groups. *p<0.05; **p<0.01; ***p<0.001. For (F), statistical significance was determined using a mixed-effects analysis followed by Tukey’s post-hoc multiple comparisons testing. A minimum of 4 male and 4 female mice per experimental group were used in all behavioral studies.

To investigate the impact of *Ficd* deletion on motor function, we evaluated mice on a five-day behavioral testing paradigm consisting of 4 days of balance beam assessments and a final day of open field testing. All animals were evaluated at 8, 12, 16, 24, 36, and 48 weeks of age (Fig. 1B). From the onset of testing, Q84/Q84; *Ficd*^+/+^ animals showed pronounced gait impairment in the 5mm balance beam traversal task. Strikingly, however, Q84/Q84; *Ficd*^-/-^ mice exhibited a near-complete rescue of motor dysfunction, performing indistinguishably from WT/WT littermates (Fig. 1C). This rescue persisted across all four days of balance beam trials and at all time-points tested, indicating that *Ficd* loss is sufficient to protect against the cumulative effects of mutant ATXN3 pathology. Q84/Q84; *Ficd*^-/-^ mice also recorded significantly fewer instances of motor abnormalities (dragging or slippage of hindlimbs) while Q84/Q84; *Ficd*^+/+^ mice exhibited increasing motor dysfunction with age (Fig. 1D). On a less sensitive motor task – traversing a 11mm round beam – Q84/Q84; *Ficd*^-/-^ mice significantly out-performed *Ficd*^+/+^ animals activity in terms of traversal time and coordination, albeit to a less-pronounced degree (Fig. S2C-D). In open field testing (30 min), *Ficd* deletion reversed hypoactivity characteristic of Q84/Q84 mice (*56*) as evidenced by an increase in total locomotor activity early (8 weeks) and late (48 weeks) in life, with a trend towards increased exploratory activity also observed at 8 weeks (Fig. 1E, Fig. S3A). Notably, the increase in total locomotor activity was attributed specifically to increased ambulatory movement in Q84/Q84; *Ficd*^-/-^ mice, while no changes in fine motor movement were detected between groups (Fig. S3B-C). Q84/Q84; *Ficd*^-/-^ mice were also observed to spend more time in the center of the open field chamber than their *Ficd*^+/+^ counterparts (Fig. S3D-E). In WT/WT littermates, *Ficd* loss had no statistically significant effect on performance in either behavior task, corroborating prior reports that *Ficd* is dispensable in the absence of physiological stress (*30*, *44*). Taken collectively, these data show that loss of FICD-mediated AMPylation robustly attenuates progressive motor dysfunction in transgenic SCA3 mice across two independent behavioral measures, suggesting a beneficial effect of decreased FICD activity on mutant ATXN3-mediated toxicity.

### Loss of FICD-mediated AMPylation decreases levels of mutant ATXN3 in affected brain regions

At 48 weeks of age, we harvested tissue from experimental animals, preserving the left hemisphere for histological studies and dissecting the right hemisphere into forebrain, brainstem, cerebellum, and cervical spinal cord for molecular analysis (Fig. 1C). To examine early-life changes associated with *Ficd* loss, we also harvested samples from an age-matched cohort of 8 week-old animals. We first assessed levels of both soluble hATXN3 and endogenous mouse ATXN3 in the cerebellum and brainstem, two primary sites affected by ATXN3 pathology in SCA3 patients. Soluble cerebellar protein lysates from young (8 weeks) animals revealed elevated hATXN3 levels in the absence of *Ficd* driven by an increase in high molecular weight (HMW) ATXN3 species (Fig. S4A-B). By 48 weeks, there was no detectable difference in monomeric ATXN3 levels between groups by western blot (Fig. 2A-B), but immunofluorescent (IF) staining for ATXN3 revealed markedly reduced nuclear ATXN3 accumulation in the deep cerebellar nuclei (DCN) of *Ficd*^-/-^ mice (Fig. S5C-D). In the *Ficd*^-/-^ brainstem, we observed reduced levels of monomeric ATXN3 early in life (Fig. S4C-D), with a pronounced reduction in HMW ATXN3 species and a trend towards overall lowering of hATXN3 noted at 48 weeks (Fig. 2C-D). IF staining in the pons, a region prominently impacted in SCA3, corroborated these findings, with little change noted at 8 weeks (Fig. S5A-B), but a substantial reduction in ATXN3 nuclear accumulation revealed in the pons of 48 week *Ficd*^-/-^ animals (Fig. 2I-J). Surprisingly, the most dramatic impact of *Ficd* loss was observed in the forebrain, where *Ficd*^-/-^ mice exhibited substantially lower levels of mutant ATXN3 compared to *Ficd*^+/+^ littermates at both young and old timepoints (Fig. S4E-F, Fig. 2E-F). In the cervical spinal cord, we found no change at 8 weeks (Fig. S4G-H), but, at 48 weeks, *Ficd*^-/-^ mice again showed markedly reduced levels of total, HMW, and human mutant ATXN3 species and increased levels of endogenous mouse ATXN3 (Fig. 2G-H). Taken collectively, these data indicate that *Ficd* loss leads to significantly lower levels of soluble human mutant ATXN3 in specific brain regions and reduced nuclear ATXN3 accumulation in aged Q84/Q84 animals, providing one potential mechanism for the observed robust behavioral rescue.

**Fig. 2.**
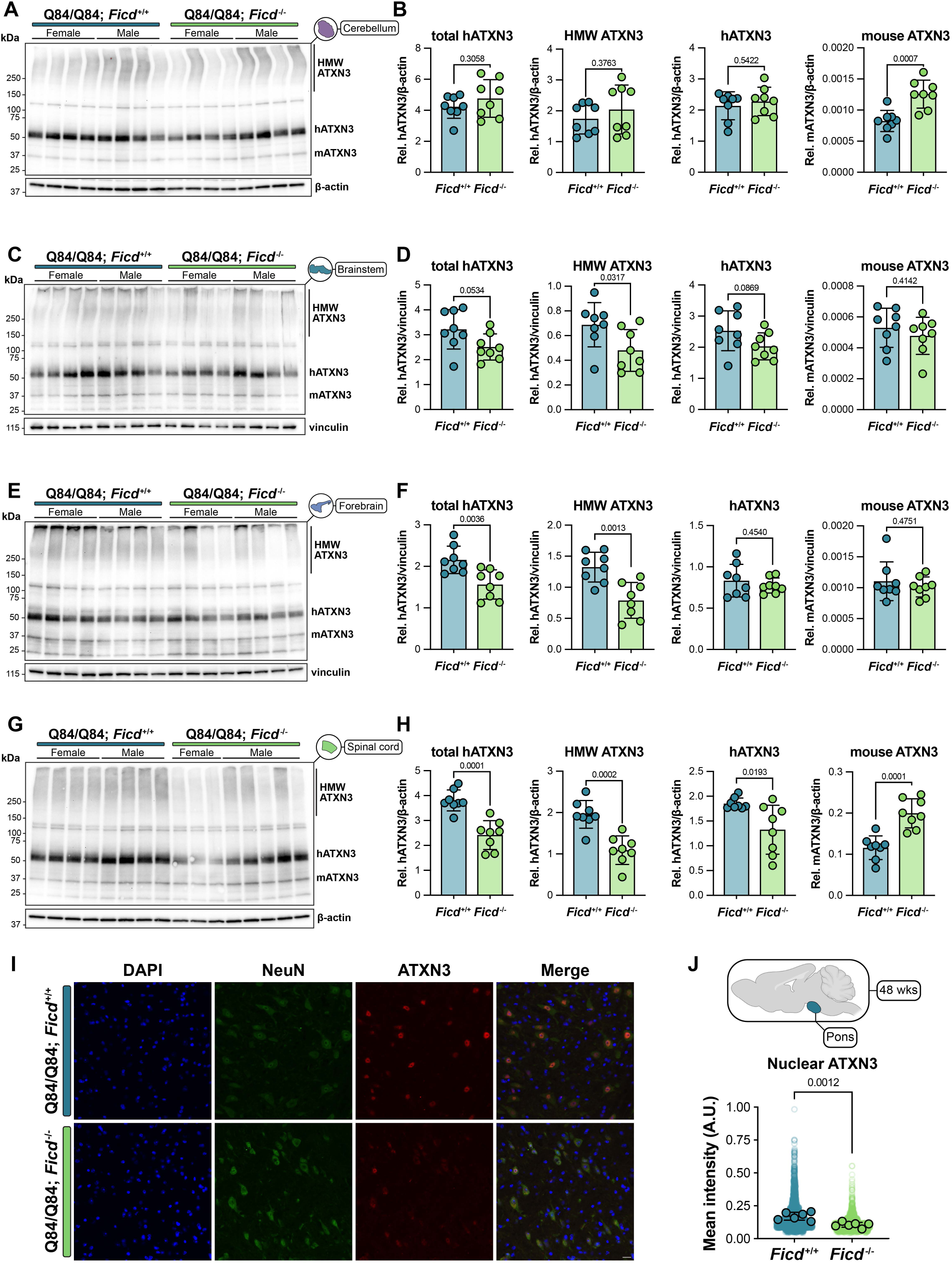
*Ficd*^-/-^ animals show decreased levels of mutant ATXN3 in affected brain regions. (A-B) Representative western blot of ATXN3 levels in the cerebellum of 48 week-old mice and quantification of total, high molecular weight (HMW), and monomeric mutant human ATXN3 (hATXN3) as well as endogenous mouse ATXN3 (mATXN3). (C-D) Representative western blot and quantification of ATXN3 levels in the brainstem of 48 week-old mice. (E-F) Representative western blot and quantification of ATXN3 levels in the forebrain of 48 week-old mice. (G-H) Representative western blot and quantification of ATXN3 levels in the spinal cord of 48 week-old mice. (I) Representative sagittal immunofluorescent images of DAPI (blue), NeuN (green), and ATXN3 (red) expression in the pons of 48 week-old mice. Scale bar = 20 μm. (J) Quantification of nuclear ATXN3 levels in NeuN+ cells (*n* = 6 images per mouse, 6 mice per genotype). Translucent data points represent individual cells, while opaque data points reflect the average intensity for each animal. Per-mouse values were used to compute significance. For (A-J), a minimum of 3 male and 3 female mice were used in all experiments. Data are mean ± SD. Statistical significance was determined using unpaired parametric *t* tests with exact p-values reported on each graph.

### The SCA3 mouse spinal cord exhibits an increased burden of AMPylated proteins

Several FICD mutations have been linked to disease in humans – most notably, Arg371Ser (*40*, *41*) and Arg374His (*42*). In both cases, the mutation occurs in the enzyme’s active site and impedes FICD’s deAMPylation abilities, resulting in a pool of AMPylated, inactive BiP. This suggests that abnormally high levels of AMPylated proteins serve as a marker of cellular stress. To this end, we assessed AMPylation levels in WT vs. SCA3 mouse brain regions by western blot using an antibody raised against threonine AMPylated BiP (Thr-AMP) (*58*). In these blots, AMPylated BiP runs at approximately ∼80 kDa and AMPylation levels are normalized to total BiP. In all regions tested, the absence of a Thr-AMP band in *Ficd*^-/-^ samples confirms the lack of endogenous AMPylation activity in *Ficd*^-/-^ mice (Fig. 3, Fig. S6). At 8 weeks, AMPylation signal was the most robust in the cerebellum, and AMPylation levels at both time-points were unchanged between WT and SCA3 groups (Fig. 3A-B, Fig. S6A-B). In the brainstem, we observed a trend towards increased AMPylation in 8 week-old female, but not male SCA3 mice, which reached significance by 48 weeks of age, reflecting possible sex-specific effects (Fig. 3C-D, Fig. S6C-D). Interestingly, at 8 weeks we observed a complete lack of AMPylation signal in forebrain and spinal cord samples across all groups, potentially explained by the hypothesis that wild-type FICD acts predominantly as a deAMPylator (*36–38*) (Fig. S6E-H). By 48 weeks of age, however, AMPylation was robustly detected in both tissues, suggesting age-dependent accumulation of AMPylated BiP may explain the appearance of Thr-AMP signal at this time-point (Fig. 3E-H). In contrast to the forebrain, where no differences were detected (Fig. 3E-F), AMPylation levels were markedly elevated in the spinal cord of both female and male SCA3 mice compared to WT littermates (Fig. 3G-H), suggesting a role for increased BiP AMPylation as a driver of SCA3 pathology in this vulnerable region.

**Fig. 3.**
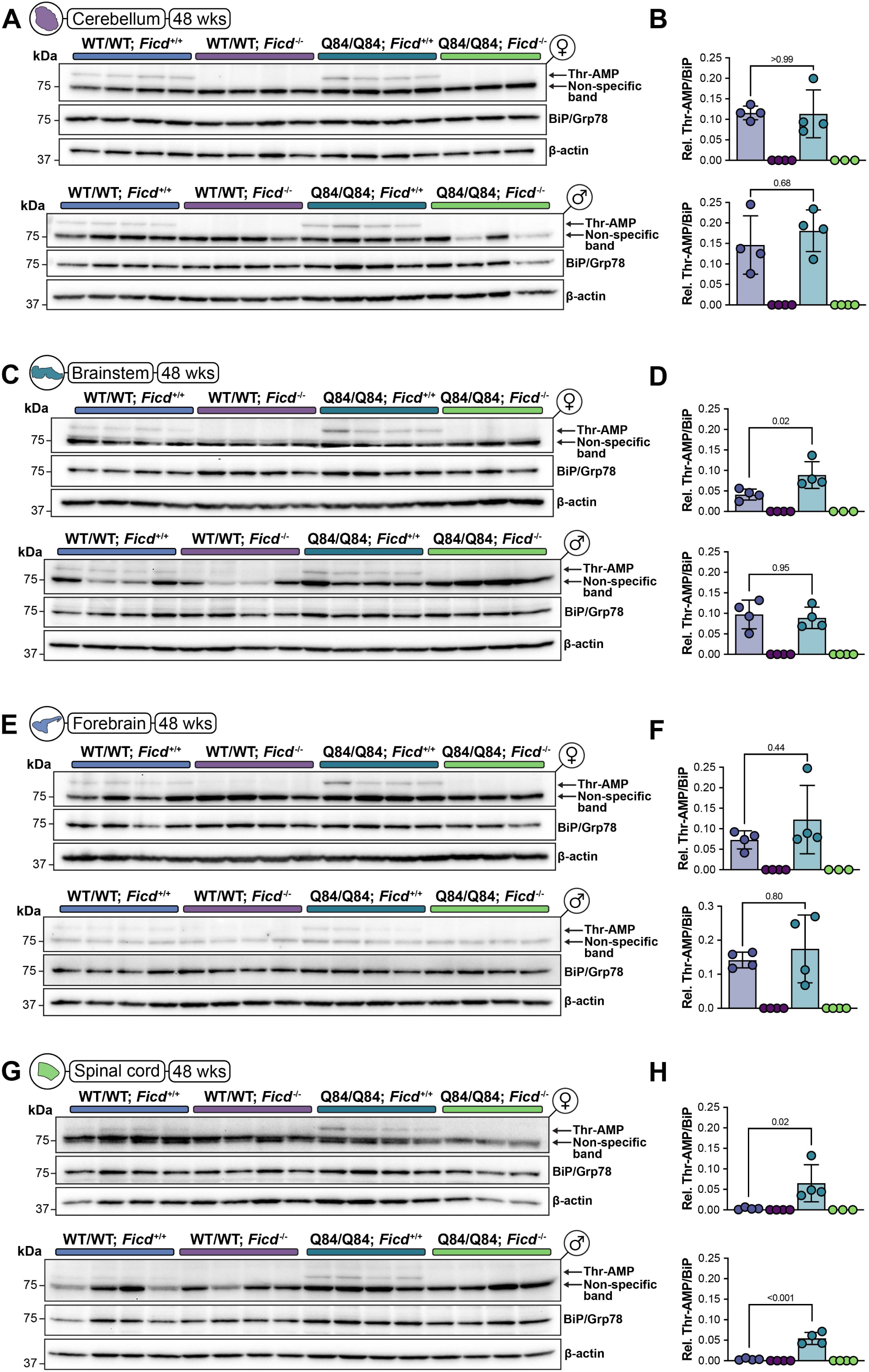
The SCA3 mouse spinal cord exhibits an increased burden of AMPylated proteins. (A-B) Western blot showing levels of AMPylated threonine (Thr-AMP), BiP, and β-actin in the cerebellum of 48 week-old female (top) and male (bottom) mice from all 4 experimental groups and quantification of AMPylated BiP levels normalized to total BiP. (C-D) Western blot and quantification of AMPylated BiP levels in the brainstem of female (top) and male (bottom) mice. (E-F) Western blot and quantification of AMPylated BiP levels in the forebrain of female (top) and male (bottom) mice. (G-H) Western blot and quantification of AMPylated BiP levels in the cervical spinal cord of female (top) and male (bottom) mice. With the exception of female Q84/Q84; *Ficd*^-/-^ mice (*n* = 3), *n* = 4 male or 4 female mice were used in all experiments. Data are shown as mean ± SD. Statistical significance was determined by one-way ANOVA with Tukey’s post-hoc multiple comparisons testing. Exact p-values are displayed on graphs.

### Unbiased proteomics reveal *Ficd* loss activates a network of myelin structural proteins, oligodendrocyte maturation factors, and lipid synthesis enzymes in the SCA3 brainstem

To further investigate the molecular changes imparted by FICD loss, we utilized data-independent mass spectrometry (DIA-MS) to compare the proteomes of all four experimental genotypes in both 8 and 48-week cohorts (Fig. 4A). At 8 weeks, principal component analysis (PCA) revealed significant overlap of *Ficd*^+/+^ and *Ficd*^-/-^ groups stratified by SCA3 transgene status (Fig. S7A), with a distinct cluster attributed to Q84/Q84; *Ficd*^-/-^ samples emerging by 48 weeks (Fig. 4B). In line with previous reports that *Ficd* loss has minimal impact on organismal physiology in the absence of stressors, we observed few differentially-expressed (DE) proteins between SCA3-*Ficd*^+/+^ and *Ficd*^-/-^ mice at either time-point (Fig. S7B, Fig. S8A) with no trends emerging in over-representation analysis (ORA). Interestingly, even fewer DE proteins were detected between SCA3^+^ *Ficd*^+/+^ and *Ficd*^-/-^ mice at 8 weeks (Fig. S7E), suggesting that the rescue is not evident at the protein level in the brainstem early in life, or is limited to changes in other regions. However, comparisons between SCA3^-^ and SCA3^+^ mice begin to show evidence of deficits in oligodendrocyte maturation and myelination at this early time point. Independent of *Ficd* genotype, SCA3^+^ animals showed a decrease in protein levels linked to myelination, axon ensheathment, and the production of lipids essential to myelin formation when compared to animals harboring the mutant *ATXN3* transgene (Fig. S7C-D, F, H). Relative to SCA3+ samples, the enrichment of cellular compartment GO terms linked to these functions, such as the myelin sheath and neurofilaments, which increase in expression during the maturation of myelinating axons (*59*), strengthen the claim that white matter dysregulation is an early feature of SCA3 pathology (Fig. S7G, I).

**Fig. 4.**
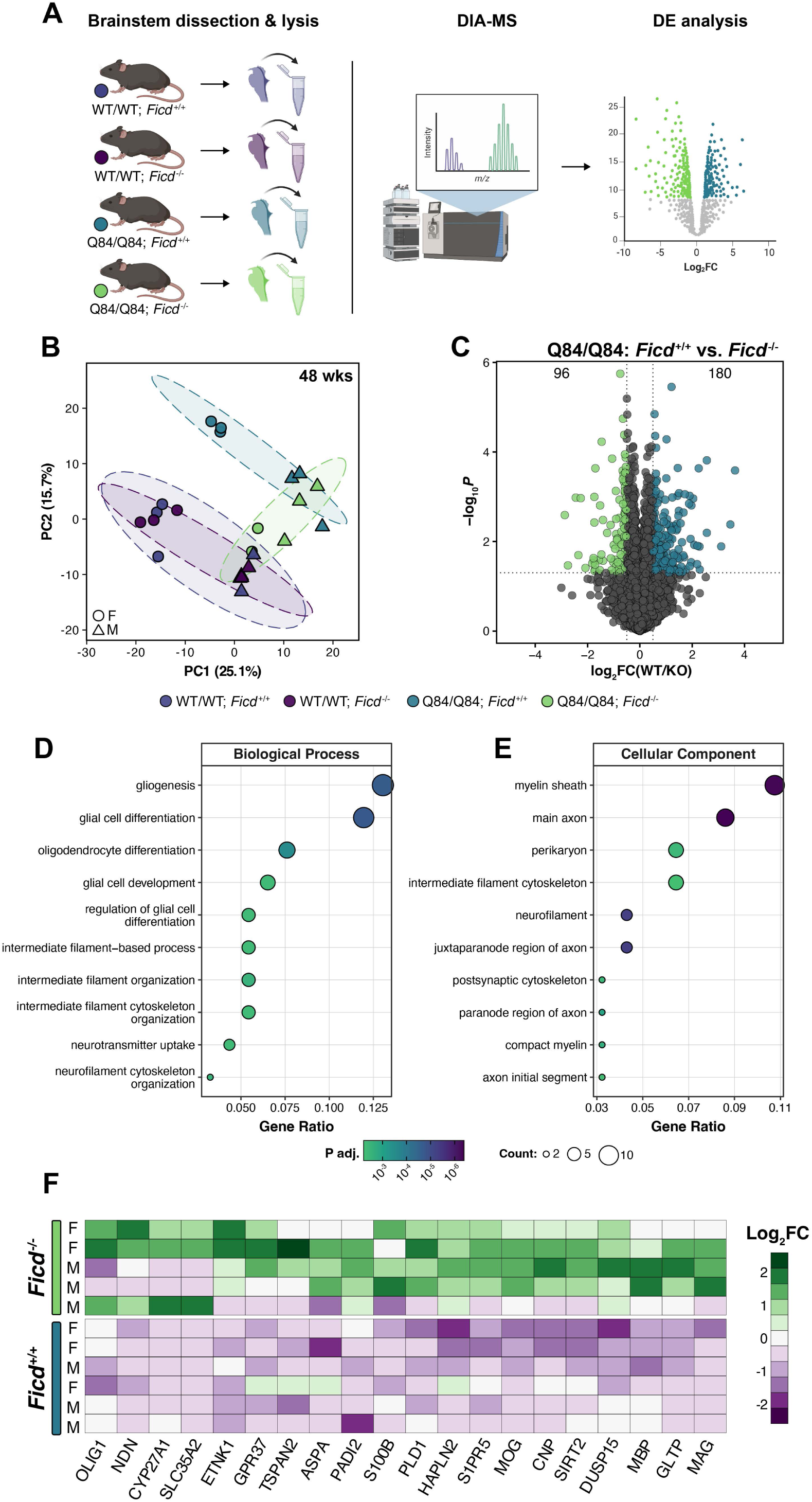
Unbiased proteomics reveal *Ficd* loss activates a network of myelin structural proteins, oligodendrocyte maturation factors, and lipid synthesis enzymes in the SCA3 brainstem. (A) Schematic depicting the experimental design employed for DIA-MS proteomics analysis of 8- and 48 week-old mouse brainstem samples. For both age cohorts, a minimum of *n* = 5 mice per genotype group were submitted for analysis. (B) Principal component analysis (PCA) plot showing clustering of 48 week-old samples by genotype. Each data point represents one mouse. Circles represent females, triangles represent males. (C) Volcano plot comparing Q84/Q84; *Ficd*^+/+^ vs. *Ficd*^-/-^ proteomes. Dashed lines reflect statistical cut-offs (log_2_FC > |0.5|, p < 0.05). Proteins upregulated in *Ficd*^-/-^ mice are colored green, and those upregulated in *Ficd*^+/+^ mice are blue. The total number of differentially expressed (DE) proteins for each group are shown in the top left and right corners. (D-E) Dot plots showing the top biological process (D) and cellular compartment (E) gene ontology (GO) terms over-represented in Q84/Q84; *Ficd*^-/-^ vs. *Ficd*^+/+^ groups. Dot size reflects the number of proteins associated with the GO term and dots are colored according to adjusted p-value (*P_adj_*). (F) Curated heat map of myelin proteins, oligodendrocyte maturation factors, and lipid synthesis enzymes upregulated in Q84/Q84; *Ficd*^-/-^ vs. *Ficd*^+/+^ mice. Each horizontal row depicts log_2_FC values from one mouse. ‘M’ and ‘F’ indicates the sex of each animal.

While few changes were observed between *Ficd*^+/+^ and *Ficd*^-/-^ Q84/Q84 mice at 8 weeks, the *Ficd* null proteome diverged significantly at 48 weeks, reflected by distinct cluster separation visualized by PCA (Fig. 4B-C). Relative to SCA3^+^; *Ficd*^+/+^ animals, ORA revealed a striking enrichment of proteins linked to gliogenesis and oligodendrocyte differentiation in the *Ficd*^-/-^ brainstem (Fig. 4D-E), suggesting that restoration of myelination deficits in the brainstem may contribute to the observed behavioral rescue.. Indeed, heatmap visualization of log_2_FC values from individual mice showed that levels of myelin structural proteins (MBP, MOG, MAG), oligodendrocyte differentiation factors (OLIG1, NDN, S1PR5), and proteins involved in lipid and glycoprotein metabolism (ASPA, PLD1, ETNK1) were markedly recovered in the *Ficd* null group (Fig. 4F). Analysis of proteomics data comparing SCA3^-^ and SCA3^+^ animals at 48 weeks continued to reflect large deficits in oligodendrocyte maturation and synthesis of myelin components in SCA3^+^ mice regardless of *Ficd* genotype, underscoring the importance of glial homeostasis in SCA3 pathology (Fig. S8B-G). Collectively, these data provide robust evidence that, in the presence of mutant ATXN3, *Ficd* deletion promotes oligodendrocyte maturation and lipid synthesis, thereby restoring glial function in the brainstem.

### *Ficd* deletion restores myelination in SCA3 mice by reversing deficits in oligodendrocyte maturation

To extend upon our proteomics findings, we next sought to confirm functional rescue of myelin structural proteins in the *Ficd*^-/-^ brainstem and expand our investigation to the spinal cord, a SCA3-vulnerable region marked by a pathological increase in AMPylation (Fig. 3G-H). Indeed, western blot analysis of 48-week brainstem lysates confirmed a pronounced rescue in levels of myelin basic protein (MBP), myelin-associated glycoprotein (MAG), and aspartoacylase (ASPA), a deacetylase essential for myelin synthesis, in Q84/Q84; *Ficd*^-/-^ animals compared to Q84/Q84; *Ficd*^+/+^ littermates (Fig. 5A-F). In the *Ficd*^-/-^ spinal cord, we similarly observed restoration of MBP levels accompanied by trends towards increased levels of MAG and ASPA in this region (Fig. 5G-L). Levels of proteolipid protein 1 (PLP1), one of the most abundant myelin structural proteins (*60*), were also rescued in the *Ficd*^-/-^ spinal cord (Fig. S9I-J) and trended towards an increase in the brainstem of *Ficd*^-/-^ mice (Fig. S9G-H). To ascertain the extent of the observed rescue, we compared normalized abundance values obtained by DIA-MS across all 4 genotypes. Compared to WT/WT controls, Q84/Q84 animals showed decreased levels of several key myelin proteins – MBP, MAG, myelin oligodendrocyte glycoprotein (MOG), PLP1, ASPA, and CNPase – with *Ficd* deletion conferring a partial rescue in all observed cases (Fig. S9A-F).In addition to proteins involved in myelin synthesis and structure, we also confirmed increased levels of the autophagic regulator, GABARAPL1, in the *Ficd*^-/-^ brainstem by western blot (Fig. S9K-L), implicating enhanced autophagic turnover as one possible explanation for lower levels of mutant ATXN3 in *Ficd*^-/-^ mice. As a whole, these findings validate our proteomics analysis and provide concrete evidence that *Ficd* deletion confers partial recovery of major myelin synthesis and structural components in the brainstem and spinal cord of SCA3 mice.

**Fig. 5.**
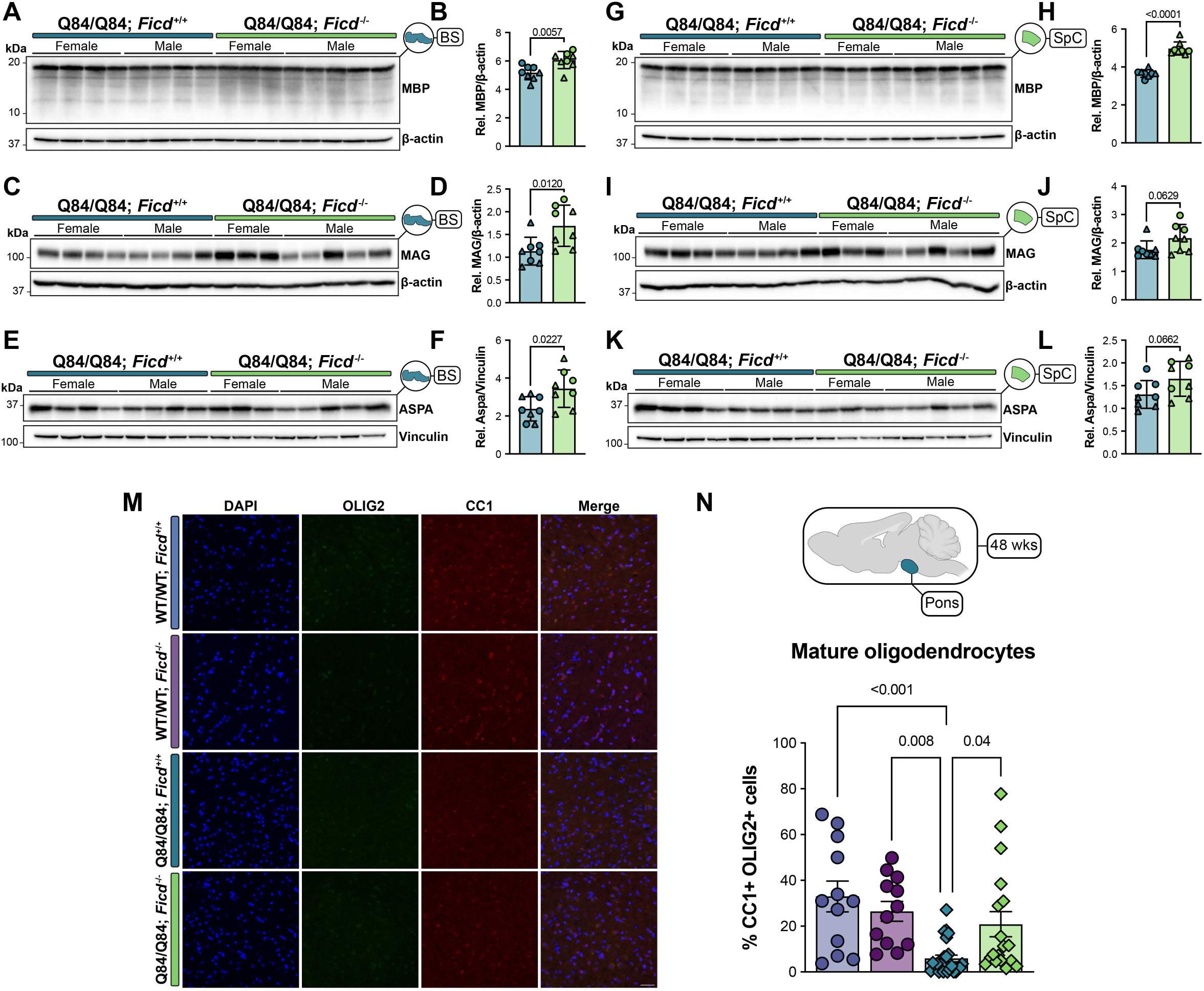
*Ficd* deletion restores myelination in SCA3 mice by reversing deficits in oligodendrocyte maturation. (A-B) Representative western blot and quantification of myelin basic protein (MBP) levels in the brainstem of 48 week-old mice. (C-D) Western blot and quantification of myelin-associated glycoprotein (MAG) levels in the brainstem of 48 week-old mice. (E-F) Western blot and quantification of aspartoacylase (ASPA) levels in the brainstem of 48 week-old mice. (G-H) Western blot and quantification of MBP levels in the cervical spinal cord. The beta-actin loading control in (G) is the same loading control shown in Fig. S9I. The same membrane was stripped and re-probed for each target. The loading control is intentionally shown in both figures for clarity. (I-J) Western blot and quantification of MAG levels in the cervical spinal cord of 48 week-old mice. (K-L) Western blot and (L) quantification of ASPA levels in the cervical spinal cord of 48 week-old mice. (M) Representative sagittal immunofluorescent images of DAPI (blue), OLIG2 (green) and CC1 (red) staining in the pons of 48 week-old mice from all four experimental groups. Scale bar = 40 μm. (N) Quantification depicting the % of CC1^+^ OLIG2^+^ double-positive cells, indicating mature oligodendrocytes (*n* = 4 images per mouse, 4 mice per genotype minimum, split evenly by sex). Data are mean ± SEM. For (A-L), *n* = 4 male and 4 female mice were analyzed per group, with the exception of female Q84/Q84; *Ficd*^-/-^ mice (*n* = 3). Data are mean ± SD. Females are circles, males are triangles. Statistical significance was determined by unpaired, parametric *t*-tests. For (M-N), a one-way ANOVA followed by Tukey’s post-hoc multiple comparisons testing was used. Exact p-values are displayed on graphs.

Previous work on glial pathology in SCA3 has linked deficits in myelin production to a pathological decrease in the number of mature, myelinating oligodendrocytes (*15*, *18*, *26*). Further, primary oligodendrocyte precursor cells (OPCs) obtained from SCA3 mice and cultured *ex vivo* exhibited the same maturation block, indicating this impairment is cell-autonomous in nature (*19*). Based on these findings and our observation that *Ficd* depletion restores myelination deficits in SCA3 mice, we asked whether loss of FICD-mediated AMPylation is sufficient to promote OPC development into mature, myelinating oligodendrocytes. To this end, we performed immunohistochemical staining in the pons for OLIG2 to mark oligodendrocyte lineage cells, and CC1, an antibody which binds QKI7, an RNA binding protein highly expressed in mature oligodendrocytes (*61*). Cells double-positive for both markers (CC1^+^ OLIG2^+^) therefore reflect myelinating oligodendrocytes. Corroborating previous findings, our analysis shows a substantial reduction in CC1^+^, OLIG2^+^ cells in Q84/Q84; *Ficd*^+/+^ animals compared to WT/WT littermate controls (Fig. 5G-H). Strikingly, however, *Ficd* loss rescues this reduction, with *Ficd* null SCA3 mice exhibiting restored levels of CC1^+^ OLIG2^+^ cells (Fig. 5G-H). Taken as a whole, these findings demonstrate that *Ficd* loss not only restores the production of key myelination factors, but also reverses a pathological block in oligodendrocyte maturation in the pons.

### Loss of FICD-mediated AMPylation rewires cholesterol biosynthesis via *Srebp2*

Having confirmed that *Ficd* deletion restores myelinating oligodendrocytes in the brainstem and cervical spinal cord of SCA3 mice, we next asked whether the loss of AMPylation supports white matter integrity by driving the synthesis of myelin-essential lipids. To this end, we performed bulk RNA-sequencing of cervical spinal cord tissue, a SCA3-vulnerable tissue that exhibits high levels of AMPylation (Fig. 6A). PCA revealed separation of all 4 groups at 48 weeks (Fig. 6B). Analyzing Q84/Q84; *Ficd*^+/+^ vs. *Ficd*^-/-^ transcriptomes, we found that gene ontology (GO) terms linked to cholesterol biosynthesis, synthesis of cholesterol intermediates, glial cell differentiation, lipid storage, and cholesterol import were significantly over-represented in SCA3 mice lacking *Ficd* (Fig. 6C-D). Cholesterol is an essential component of myelin, comprising ∼40% of its total lipid content (*62*), and is synthesized *de novo* in the CNS via the isoprenoid biosynthetic pathway (*63*). Cholesterol synthesis is regulated by the sterol regulatory element-binding protein 2 (SREBP2). Normally found in the ER, SREBP2 is transported to the Golgi apparatus when cholesterol levels are low, where it is cleaved into its active form, translocates to the nucleus, and binds sterol regulatory elements (SRE) to activate cholesterol synthesis (*64*, *65*). Given that SREBP2 and FICD are ER-resident proteins, and Grp78/BiP inhibition reduces SREBP2 levels (*66*), we hypothesized that *Ficd* loss promotes cholesterol synthesis via SREBP2 due to an increased pool of unAMPylated BiP. We first confirmed upregulation of cholesterol synthesis genes in *Ficd*^-/-^ SCA3 mice by qPCR. Indeed, Q84/Q84; *Ficd*^-/-^ mouse spinal cord samples showed increased transcription of several enzymes involved in cholesterol synthesis (*Hmgcr*, *Hmgcs1*, *Sqle*) as well as *Srebf2* (the gene encoding SREBP2) upregulation (Fig. 6E-I). Profiling of transcriptomic changes across experiments revealed a marked downregulation of sterol biosynthesis genes in Q84/Q84; *Ficd*^+/+^ mice relative to WT/WT controls that was attenuated in the absence of *Ficd*, confirming the biological relevance of this partial rescue (Fig. S10A). We also observed this same increase in the brainstem, but not the cerebellum, of *Ficd*^-/-^ mice, suggesting a region- or age-dependent effect (Fig. S10B-K). To determine if *Ficd* loss influences cholesterol synthesis via SREBP2, we assessed levels of the pro-(SREBP2-P) and active (SREBP2-N) forms of SREBP2 by western blot. In the spinal cord, Q84/Q84 *Ficd*^-/-^ mice exhibited moderately elevated levels of both pro- and nuclear SREBP2, consistent with activation of SREBP2-dependent gene expression alongside increased levels of SREBP2 transcripts (Fig. 6J-M). In contrast, we did not observe this effect in the Q84/Q84 *Ficd*^-/-^ brainstem (Fig. S10L-O). From this, we conclude that the loss of FICD-mediated AMPylation may promote cholesterol production, in part, through SREBP2-dependent gene expression.

**Fig. 6.**
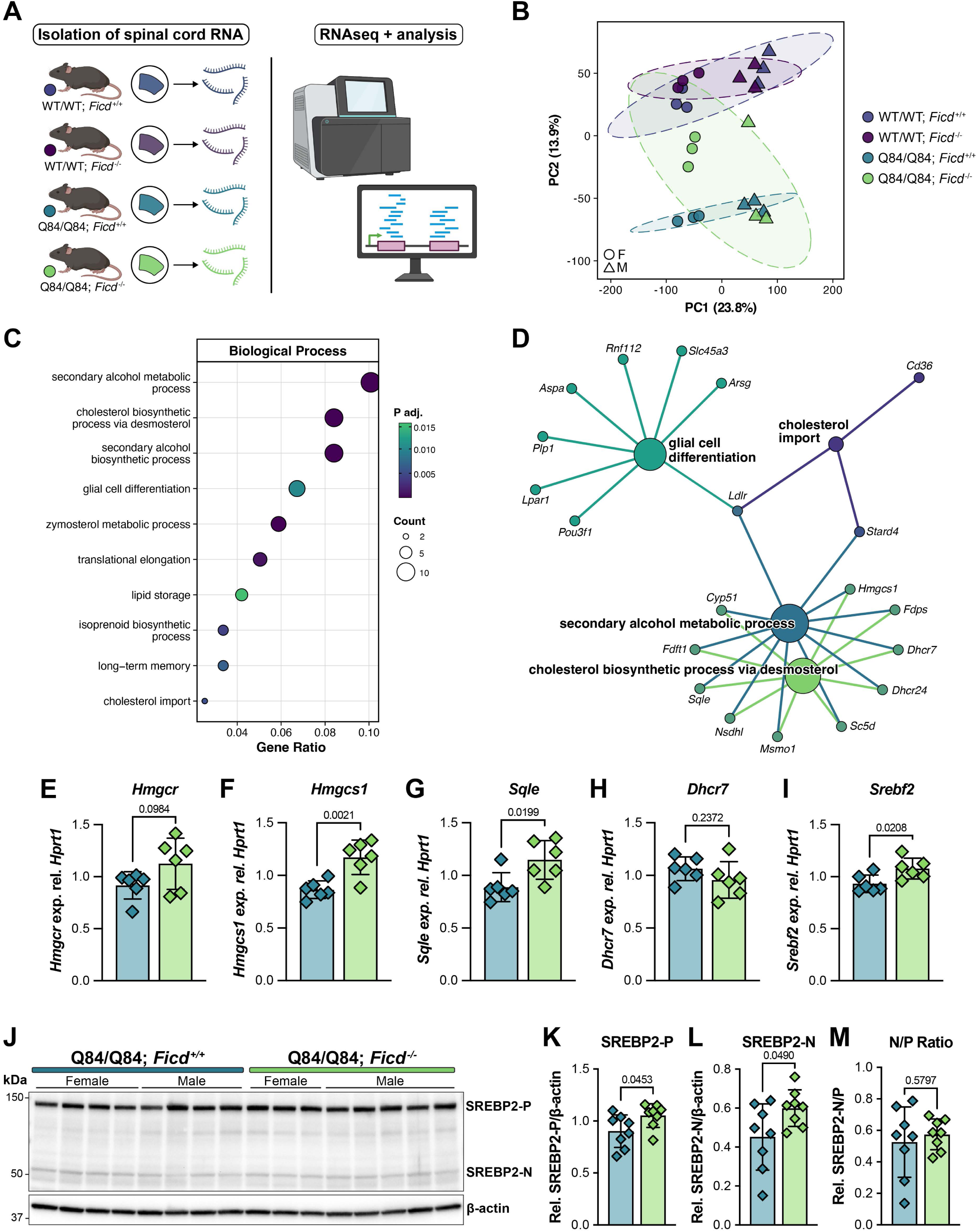
Loss of FICD-mediated AMPylation rewires cholesterol biosynthesis via SREBP2. (A) Schematic describing the experimental design employed for bulk RNA-seq analysis of 48-week mouse cervical spinal cord tissue (*n* = 6 mice per genotype, split evenly by sex). (B) Principal component analysis plot showing separation and clustering of transcriptomic data by genotype. Female samples are circles, males are triangles. (C) Dot plot of gene ontology (GO) terms significantly over-represented in Q84/Q84; *Ficd*^-/-^ vs. *Ficd*^+/+^ spinal cord tissue. Dot size reflects the number of genes associated with the GO term and dots are colored according to adjusted p-value (*P_adj_*). (D) Gene-concept (“cnet”) plot depicting the genes associated with representative GO terms identified in (C) and their functional relationships. Colored lines and nodes denote linkage to GO terms and overlap between categories. (E-I) Quantitative PCR (qPCR) confirming significantly increased expression of cholesterol synthesis genes (E) *Hmgcr*, (F) *Hmgcs1*, (G) *Sqle*, (H) *Dhcr7*, and (I) *Srebf2* in the spinal cord of Q84/Q84; *Ficd* null mice at 48 weeks (*n* = 6 mice per genotype, split evenly by sex). Data are mean ± SD. (J) Representative western blot showing levels of the precursor (-P) and spliced, nuclear (- N) forms of SREBP2 in the spinal cord of 48 week-old mice (*Ficd*^+/+^ *n* = 4 male, 4 female mice; *Ficd*^-/-^ *n* = 3 female, 5 male mice). (K-M) Quantification of (K) precursor (SREBP2-P), (L) nuclear (SREBP2-N), and (M) ratio of nuclear to precursor SREBP2. Data are mean ± SD. For (E-I, K-M) statistical significance was determined by unpaired, parametric *t* tests. Exact p-values displayed on graphs.

## DISCUSSION

In recent years, glial cell involvement has become increasingly recognized as a feature of SCA3 and polyglutamine disease pathology more broadly (*15*, *67–69*). While earlier studies identified oligodendrocyte involvement late in disease (*13*, *14*, *16*, *25*), additional preclinical work has since characterized deficits in oligodendrocyte maturation as a pathological feature evident early in the disease process. The translational relevance of these findings has since been bolstered by neuroimaging studies which highlight white matter changes as an early feature of disease in SCA3 patients (*23*, *70*, *71*). Importantly, evidence that ATXN3 loss does not impact oligodendrocyte function (*14*) combined with studies implicating impaired oligodendrocyte maturation as a driver of pathology in Huntington’s disease (*67*, *68*) suggest that oligodendrocyte dysfunction may indeed be a shared pathological mechanism imparted by gain-of-function toxicity arising from the aberrant CAG repeat expansions which define these disorders. This potential for a shared pathogenic mechanism that may be targeted for therapeutic development has underscored the need for further research into the drivers of oligodendrocyte dysfunction.

In this study, we explore how loss of *Ficd* alters pathology in a SCA3 mouse model. This bifunctional enzyme catalyzes both the addition and removal of adenosine monophosphate (AMP) to target proteins, allowing for temporal regulation of its substrates. *Ficd* is most well characterized as a regulator of the ER-resident, HSP70 family chaperone, BiP, though additional AMPylation targets have since been identified (*72*, *73*). Mounting evidence has implicated FICD-mediated AMPylation as a key regulator of ER stress (*34*, *45*, *46*, *74*), suggesting its involvement in heavily ER-dependent processes. Several studies indicate a role for AMPylation in the pathology of neurodegenerative diseases such as Parkinson’s disease (*49*, *50*, *73*). More recently, we showed that loss of AMPylation suppresses polyglutamine toxicity in *C. elegans* (*51*) through a UPR^ER^-dependent signaling mechanism. Still, how *Ficd* activity alters toxicity in polyglutamine diseases has remained poorly understood.

Our data indicate that constitutive *Ficd* deletion is sufficient to rescue motor function in mice expressing full-length mutant human ATXN3 even in late stage disease. Post-mortem analysis of 48 week old mice showed that *Ficd* null animals exhibited lower levels of neuronal nuclear ATXN3 accumulation, suggesting one explanation for the observed rescue. However, the mechanism(s) by which *Ficd* loss clears mutant ATXN3 in our model are unclear. ATXN3 is predominantly a cytosolic and nuclear protein (*75*), while *Ficd* is thought to reside in the ER. We have previously reported that loss of the *C. elegans* ortholog, *fic-1*, activates expression of genes linked to ER-associated degradation (ERAD) in the context of polyQ aggregation (*51*). As ATXN3 has been shown to interact with VCP/p97 at the cytosolic face of the ER lumen (*76*), one possibility is that *Ficd* loss incites ATXN3 clearance via ERAD. Other studies have implicated *Ficd* in the regulation of autophagy either through the direct modification of lysosomal cathepsins (*72*) or indirect activation of protein clearance mechanisms through hormetic UPR^ER^ induction (*47*). Our analysis of proteomic data from *Ficd*^-/-^ mouse brainstem samples revealed upregulation of cathepsin D (CTSD) as well as the autophagic flux regulator GABARAPL1 (*77*, *78*), suggesting the possibility that *Ficd* deletion activates autophagic degradation processes. Still, targeted experiments are needed to pinpoint the exact cause of reduced ATXN3 nuclear accumulation and changes in ATXN3 levels observed in *Ficd* null mice.

Perhaps our most striking molecular finding is that *Ficd* deletion restores deficits in oligodendrocyte maturation in SCA3 mice. Myelin production is a metabolically-demanding task and endoplasmic reticulum stress in oligodendrocytes is widely accepted as a driver of various myelinopathies (*29*, *79–82*). Similarly, reduced cholesterol levels have been implicated previously in both Huntington’s disease (*83*) and SCA3 (*84*), with one study reporting that over-expression of a rate-limiting cholesterol synthesis enzyme was sufficient to rescue SCA3 mice (*85*). Here, we provide compelling evidence linking the loss of FICD-mediated AMPylation to the promotion of oligodendrocyte differentiation and the recovery of myelin proteins depleted in SCA3 models. Previous reports demonstrate that the major AMPylation target, Grp78/BiP, is essential for oligodendrocyte survival and its inactivation leads to the loss of myelinating oligodendrocytes (*52*). In this study, we find increased levels of AMPylated BiP in the spinal cord of late-stage SCA3 animals. This suggests that AMPylation-induced BiP inactivation could contribute to cellular dysfunction in this model.. Based on these studies and our reported findings, we propose that *Ficd* deficiency ameliorates motor phenotypes in SCA3 by preventing the pathological sequestration of inactivated BiP induced by AMPylation, boosting cellular proteostasis to lower levels of mutant ATXN3 and promote oligodendrocyte maturation. Future work aimed at deciphering the cell type-specific effects of *Ficd* deletion, such as through conditional *Ficd* knock-out strains (*45*), will be essential for identifying the cell types and signaling pathways impacted by *Ficd* loss. Taken as a whole, our results expand upon our previous findings that loss of FICD-mediated AMPylation suppresses polyglutamine toxicity and characterize *Ficd* as a novel regulator of myelination and oligodendrocyte maturation in the context of SCA3. Additional studies capitalizing on recently reported small molecule FICD inhibitors (*86*) may hold promise for the development of therapies alleviating oligodendrocyte dysfunction in SCA3 and related diseases.

### Limitations of this study

In this study, we use a constitutive full-body *Ficd* deletion mouse model, which prevents us from drawing conclusions regarding how *Ficd* loss in neurons and oligodendrocytes only, along with putative non-cell autonomous consequences, may contribute to the observed rescue. Given FICD’s near-ubiquitous expression in the brain (*87*) future studies should employ conditional *Ficd* deletion models in conjunction with cell type-specific driver lines to address this question. Similarly, we utilize bulk RNA sequencing analysis of the mouse spinal cord to identify cholesterol biosynthesis as one pathway promoted by the absence of *<u>Ficd</u>*. While oligodendrocytes are the most ubiquitous cell type in the spinal cord (*88*) and peripheral oligodendrocytes produce cholesterol at higher levels than their CNS counterparts (*89*), we cannot exclude the involvement of other cell types, such as astrocytes, in this finding. Going forward, single-cell transcriptomic analysis would be best suited to disentangle these results. Informed by our behavioral and motor rescue data, we chose to analyze 8- and 48-week SCA3 animals to capture both early symptomatic and late-stage disease. However, our molecular analyses in 8 week samples provided limited insight into the observed motor rescue. As we could not characterize samples from mid-stage (e.g. 16- or 24-weeks) SCA3 animals, we are unable to determine at what point in the disease process significant molecular changes from *Ficd* loss appear. Lastly, we focus on BiP AMPylation as a primary read-out for FICD activity. While BiP is the best characterized and most clinically-relevant AMPylation target, more recent studies have reported additional FICD substrates (*72*, *73*). As such, we cannot exclude that targets beyond BiP may play a role in the beneficial effects of *Ficd* deletion reported here.

## MATERIALS AND METHODS

### Animals

All animal procedures were approved by the University of Michigan Institutional Animal Care and Use Committee (IACUC) and conducted in accordance with the United States Health Service’s Policy on Humane Care and Use of Laboratory Animals. Mice were housed in ventilated cages and maintained on a standard 12-hour light/dark cycle, with standard chow and water *ad libitum*. The YAC-Q84 mouse model (*55*), which carries a transgene expressing the full-length human *ATXN3* gene, was used as a model for spinocerebellar ataxia type 3 (SCA3). To study the impact of *Ficd* deletion, we crossed hemizygous YAC-Q84 mice with *Ficd^-/-^* null animals (*44*) and established a colony using breeders that were hemizygous for the YAC transgene and heterozygous for the *Ficd* null allele (Q84/WT; *Ficd^+/-^*) (Fig. 1A). Both lines were maintained on a C57BL/6 background.

Genotyping was performed using DNA extracted from tail biopsies collected prior to weaning and was confirmed post-mortem. Presence/absence of the YAC-Q84 transgene and *Ficd* genotype status were determined by standard PCR as described previously (*44*, *90*). Reactions were performed with GoTaq Green Master Mix (Promega) using a ProFlex PCR System (Applied Biosystems). All PCR primers used in this study are listed in Supplementary Table S1. YAC-Q84 mouse hemi-vs. homozygosity was determined via qPCR by amplification of a fragment of the *ATXN3* transgene (Assay ID: AP7DT7R, ThermoFisher) and normalizing to a genomic fragment of the mouse *Actb* gene (Assay ID: Mm02619580_g1, ThermoFisher). The absence of *Ficd* gene expression in *Ficd*^-/-^ animals was confirmed by qPCR performed on brainstem RNA obtained post mortem (Fig. S1A-B). All qPCR reactions were performed with either PowerUp SYBR Green Master Mix (Applied Biosystems) or TaqMan Fast Advanced Master Mix (Applied Biosystems) and ran on a StepOnePlus Real-Time PCR System (Applied Biosystems). The *ATXN3* CAG trinucleotide repeat lengths of homozygous Q84 mice were determined by short tandem repeat (STR) analysis (Transnetyx) using primers for *ATXN3*. The mean ± SD repeat size for the 8 week-old cohort was 75.28 ± 2.35 for Q84/Q84; *Ficd^+/+^* animals and 75.79 ± 3.09 for Q84/Q84; *Ficd^-/-^* animals. For the 48 week cohort, mean ± SD repeat size was 73.68 ± 2.68 for Q84/Q84; *Ficd^+/+^* animals and 75.63 ± 2.4 for Q84/Q84; *Ficd^-/-^* animals. Identification numbers and CAG repeat sizes for all experimental animals (where applicable) are listed individually in Supplementary Tables S2 and S3.

### Transcardiac perfusion and tissue processing

At 8 or 48 weeks of age, animals were anesthetized via intraperitoneal injection of a ketamine/xylazine cocktail (100 mg/kg ketamine per 10 mg/kg xylazine) and perfused transcardially with PBS. The brain was removed, cut into two hemispheres, and the right hemisphere was macro-dissected into cerebellum, brainstem, forebrain, and cervical spinal cord for RNA and protein extraction (Fig. 1C). Samples were immediately placed on dry ice and subsequently stored at -80 °C. For histological studies, the left hemisphere was post-fixed overnight in 4% PFA (Electron Microscopy Sciences), followed by 30% sucrose, and frozen with chilled 2-methylbutane for storage at -80°C. Macro-dissected brain and cervical spinal cord samples were homogenized in PBS containing protease and phosphatase inhibitors (Thermo Fisher Scientific) using a bead mill (Bullet Blender, Next Advance), with an aliquot of homogenate reserved at -80 °C for RNA extraction. The remaining homogenate was suspended in chilled RIPA buffer and sonicated at 50% amplitude on ice in 30 sec pulse intervals until clear. Protein lysates were cleared by centrifugation in a chilled benchtop centrifuge for 30 min at 16,100 x g and stored at -80 °C.

### Motor evaluation

*Ficd* wild-type (*Ficd^+/+^*) or *Ficd* knock-out (*Ficd^-/-^*) mice homozygous or negative for the YAC transgene (Q84/Q84, WT/WT) were assessed in longitudinal motor function testing starting at 8 weeks and subsequently at 12, 16, 24, 36, and 48 weeks of age (± 1 week) (Fig. 1B). Testing groups were comprised of littermate male and female mice. Weights were recorded on the first day of each 5-day evaluation period, consisting of 4 days of balance beam and one day of open field testing. All testing was carried out by the same experimenter, in the same room, at approximately the same time of day. Animals were permitted to habituate to the testing environment for at least 15 minutes prior to assessment. Motor coordination and balance were tested by assessing the ability of animals to traverse a plexiglass beam (44 cm long, 53 cm high) from a clear platform (20 x 20 cm) to an enclosed black box (20 x 20 x 20 cm). Mice were tested on both an 11 mm round beam and a 5 mm square beam, crossing for two trials per beam on each of 4 testing days. Time to traverse the beam was recorded for each trial with a maximum cut-off of 20 sec per run. Instances of hindlimb dragging or slipping were also scored for each trial. Locomotor and exploratory activities were assessed by placing animals in a photobeam activity system open field apparatus (San Diego Instruments) for 30 minutes. Mouse total locomotor activity corresponds to the total number of photobeam breaks, while exploratory activity was measured by the number of rears.

### Western blot

Protein concentrations were determined by BCA assay (Pierce, Thermo Fisher Scientific). The amount of protein loaded was optimized for target expression and ranged between 10-50 μg per lane. Samples were resolved using SDS-PAGE on either fixed-percentage (10 or 15%) or 4-20% gradient gels (Bio-Rad, #5671094) and transferred to 0.2 μm PVDF membranes overnight at 4 °C. Membranes were incubated overnight at 4 °C with primary antibodies diluted in TBS-T containing 5% BSA (Dot Scientific, #DSA30075). Protein bands were visualized by incubation with HRP-conjugated anti-mouse or anti-rabbit secondary antibodies (1:10,000; Invitrogen, #A16078, #A16110) followed by treatment with Immobilon ECL reagent (Millipore-Sigma, #WBKLS0500) and exposure on an iBright 1500 imager (Invitrogen). Band intensities were quantified using Fiji/ImageJ (National Institutes of Health) (*91*). Supplementary Table S4 lists all primary antibodies used for western blots in this study.

### Protein digestion for DIA-MS

Proteins were extracted from macro-dissected brainstem samples and quantified as described above. Samples (50 μg each) were submitted to the Proteomics Resource Facility at the University of Michigan for processing and mass spectrometry data acquisition. Briefly, upon reduction (5 mM DTT, for 30 min at 45 °C) and alkylation (15 mM 2-chloroacetamide, for 30 min at room temperature) of cysteines in samples, the proteins were precipitated by adding 6 volumes of ice-cold acetone followed by overnight incubation at -20 °C. The precipitate was spun down, and the pellet was allowed to air dry. The pellet was resuspended in 0.1M TEAB and digested overnight (∼16 hrs) with trypsin/Lys-C mix (1:40 protease:protein; Promega) at 37 °C with constant mixing using a thermomixer. Digestion was stopped by adding formic acid to a final 5% (v/v) and desalted using SepPak C18 cartridges according to manufacturer’s protocol (Waters). Eluted peptides were dried in a vacufuge and stored at -80 °C until MS analysis.

### DIA-MS analysis using Orbitrap Ascend

An Orbitrap Ascend Tribrid equipped with FAIMS source (Thermo Fisher Scientific) and a Vanquish Neo UHPLC were used to acquire the data. Approximately 1 μg of the sample was resolved on an Easy-Spray PepMap Neo column (75 μm i.d. x 50 cm; Thermo Scientific) at the flow-rate of 300 nl/min using 0.1% formic acid/90% acetonitrile gradient system (3-19% acetonitrile in 72 min; 19-29% acetonitrile in 28 min; 29-41% in 20 min followed by 10 min column wash at 95% acetonitrile and re-equilibration) and directly sprayed onto the mass spectrometer using EasySpray source (Thermo Fisher Scientific). The FAIMS device was operated in standard resolution mode with a nitrogen gas flow of 4.2 L/min, electrode temperatures of 100 °C, and a dispersion voltage of -5000 V. A single compensation voltage (CVs) of -50 was employed to select ions that enter the mass spectrometer for MS1 scan and MS/MS cycles. The mass spectrometer was set to collect MS1 scan (Orbitrap; 350-1650 m/z; 120K resolution; AGC target of 250%; max IT of 50 ms). DIA-MS (tMSn) data was collected with 24 variable-width windows overlapping by 1 Da. Exact window widths used for data collection are listed in Supplementary Table S5. The DIA-MS fragmentation parameters were: 32% NCE; 30K resolution; 100% AGC target; 59 ms max IT.

### DIA-MS proteomics data analysis

DIA-MS data from the young (8 week) and aged (48 week) cohorts were analyzed separately. For each cohort, the raw files were converted to mzML format using MSConvert from ProteinWizard and analyzed with the FragPipe computational platform (v23.1) using the default DIA_SpecLib_Quant workflow. MSFragger (v4.3) (*92*, *93*) in DIA mode was run to search the data against the UniProt mouse reference proteome (UP000000589; downloaded 2025-08-11), appended with an equal number of reserved decoy sequences and common contaminants. Precursor and fragment mass tolerances were set to 20 ppm, trypsin was specified as the enzyme with up to two missed cleavages allowed, and peptide lengths were restricted to 7–50 amino acids. Oxidation of methionine and protein N-terminal acetylation were specified as variable modifications, and carbamidomethylation of cysteine was set as a fixed modification

The search results were further processed using MSBooster (*94*) for deep learning-based rescoring, followed by Percolator (*95*) for PSM validation, ProteinProphet (*96*) for protein inference, and Philosopher (*97*) for false discovery rate (FDR) control at 1% FDR at the PSM, peptide, and protein levels. Spectral libraries were then generated with EasyPQP and used by DIA-NN (*98*) to extract and quantify precursors and proteins from the DIA data. The resulting report.pg_matrix.tsv files from each cohort were used for downstream analysis. Differential protein analysis was performed using Limma’s moderated t-test via FragPipe-Analyst (*99*). Proteins with an absolute fold change ≥|0.5| and a p-value <0.05 were considered differentially expressed.

### Immunohistochemistry

PFA-fixed, cryoprotected left hemispheres were embedded in OCT compound and cut using a cryostat to collect a series of 10 μm sagittal sections mounted on glass slides and stored at -80°C until use. For immunofluorescence studies, slides were washed 3 times in 1x PBS and incubated for 20 min at 100 °C in 0.01 M sodium citrate buffer (pH 8.5) for antigen retrieval (*100*). After cooling, slides were again washed in PBS and blocked for 1 hr at room temperature in M.O.M. Mouse Ig Blocking Reagent (Vector Laboratories, BMK-2202) diluted in PBS with 5% normal goat serum according to the manufacturer’s instructions. After blocking, slides were washed twice in PBS and incubated for 15 min in M.O.M. diluent in PBS prior to incubation overnight in primary antibodies at 4 °C. Primary antibodies used for immunofluorescence in this study include mouse anti-ATXN3 (1H9) (1:500, MAB5360, Millipore), rabbit anti-NeuN (1:500, 26975-1-AP, Proteintech), mouse anti-APC (CC-1) (1:500, OP80, Millipore), and rabbit anti-OLIG2 (1:500, P21954, Invitrogen). The following day, slides were washed 3 times in PBS and incubated with Alexa Fluor Plus 488 goat anti-mouse or anti-rabbit (1:1000, A32723 or A32731, Invitrogen) and Alexa Fluor Plus 568 goat anti-mouse or anti-rabbit (1:1000, A11031 or A11011, Invitrogen) secondary antibodies diluted in PBS with 5% normal goat serum in the dark at room temperature for 1 hr. All slides stained with DAPI (Sigma) for 15 min at room temperature and mounted with VECTASHIELD HardSet Antifade Mounting Medium (Vector Labs). Imaging was performed using a Nikon Ti2 widefield microscope at 20x resolution in the pons and deep cerebellar nuclei (DCN). Anatomical regions of interest (ROIs) were extracted from larger 20x stitched scans using Fiji (*91*) and the resulting images were analyzed using Cellpose (v3.1.1.2) (*101*) and CellProfiler (v4.2.8) (*102*, *103*).

### RNA isolation and qRT-PCR

Mouse brain and spinal cord tissue crudely homogenized in PBS was suspended in 800 μL TRI Reagent (Zymo Research, R2071) and lysed in a TissueLyser III Bead Mill (Qiagen). Depending on the amount of input tissue, RNA isolation was performed using either the Direct-Zol RNA Miniprep Plus Kit or the Direct-Zol RNA Microprep Kit (Zymo Research, R2071 or R2060) according to manufacturer’s instructions. RNA was eluted in RNase-free water and assessed by Nanodrop to determine sample concentration and purity. For qRT-PCR (qPCR) analysis, 0.25-1 μg of total RNA was reverse-transcribed using the High-Capacity cDNA Reverse Transcription Kit (Applied Biosystems, #4368814). qPCR reactions were prepared using PowerUp SYBR Green Master Mix (Applied Biosystems, A25742) with 5-10 ng of input cDNA used per reaction. All reactions were plated in triplicate on 384 well plates and ran on a QuantStudio 6 Pro (Applied Biosystems). Data were normalized to the reference gene *Hrpt1*, which was selected from a screen of 4 candidate genes as having the highest comprehensive stability ranking as determined by RefFinder (*104*). Relative gene expression was calculated using the comparative Ct (2^-ΔΔCT^) method. Supplementary Table S6 lists all RT-qPCR primers used in this study.

### RNA sequencing and differential expression analysis

Purified total RNA from mouse cervical spinal cord (300 ng/sample) was submitted to Plasmidsaurus for bulk RNA sequencing using Illumina Sequencing Technology. Downstream statistical analysis on the resulting gene-level count matrix was conducted independently in R (v4.5.2) using edgeR (Bioconductor, v4.8.2) (*105*) to identify differentially expressed genes. Gene ontology (GO) over-representation analysis (ORA) was performed using the clusterProfiler package (Bioconductor, v4.18.4) (*106*) and the *org.Mm.eg.db* mouse genome annotation database (Bioconductor, v3.22.0). Gene-concept network (“cnet”) plots were generated using the enrichplot package (Bioconductor, v1.32.0).

### Statistical analysis, graphics, and illustrations

All statistical tests were performed in GraphPad Prism (v11.0.0) unless noted otherwise. The specific statistical tests used are described in the legends for each figure panel. A p-value of p < 0.05 was used as a cut-off to determine statistical significance. Graphics were created in Adobe Illustrator (v30.5.1) and BioRender.

### Generative AI use disclosure statement

The generative AI models Gemini 3.1 Pro and Gemini 3.5 Flash (Google DeepMind, 2026) were used solely to assist in coding for the visualization of proteomic and transcriptomic data presented in this study.

## Acknowledgments

We thank the members of the Truttmann lab for helpful comments and discussion and Anna J. Barget from the Costa lab for assistance with mouse work. We would also like to thank Hayley S. McLoughlin for sharing the anti-MBP antibody, the UM Proteomics Research Facility for DIA-MS data acquisition, and the UM Microscopy Core for equipment access.

## Funding

National Institutes of Health grant T32GM007315-43 (K.V.P.)

National Institutes of Health grant F31NS127485 (K.V.P.)

National Institutes of Health grant R35GM142561 (M.C.T.)

National Institutes of Health grant R35NS122302 (H.L.P.)

National Ataxia Foundation (M.C.T.)

UM Paul F. Glenn Center for Biology of Aging Research (M.C.T.)

National Ataxia Foundation (M.d.C.C.)

Michigan Medicine Pandemic Research Recovery – Hardship Award (M.d.C.C.)

National Institutes of Health grant SBIR-R44NS127711-01 (M.d.C.C.)

Cayman Biomedical Research Institute (M.d.C.C.)

## Author contributions

Conceptualization: K.V.P., M.d.C.C., M.C.T.

Data curation: K.V.P., Y.D.

Formal analysis: K.V.P., Y.D.

Funding acquisition: K.V.P., H.L.P., M.d.C.C., M.C.T.

Investigation: K.V.P.

Methodology: K.V.P., M.d.C.C., M.C.T.

Project administration: K.V.P., M.d.C.C., M.C.T.

Resources: K.V.P., A.I.N., M.d.C.C., H.L.P., M.C.T.

Software: K.V.P., Y.D., A.I.N.

Supervision: K.V.P., M.d.C.C., M.C.T.

Validation: K.V.P., Y.D.

Writing – original draft: K.V.P.

Writing – review & editing: K.V.P., M.d.C.C., H.L.P., M.C.T.

## Data, code, and materials availability

DIA-MS and RNAseq data generated in this study will be made available in an approved online repository by the time of publication. All tabulated data and code used to generate the figures presented in this study will be deposited on Zenodo by the time of publication.

## Supplementary materials

**Fig. S1.**
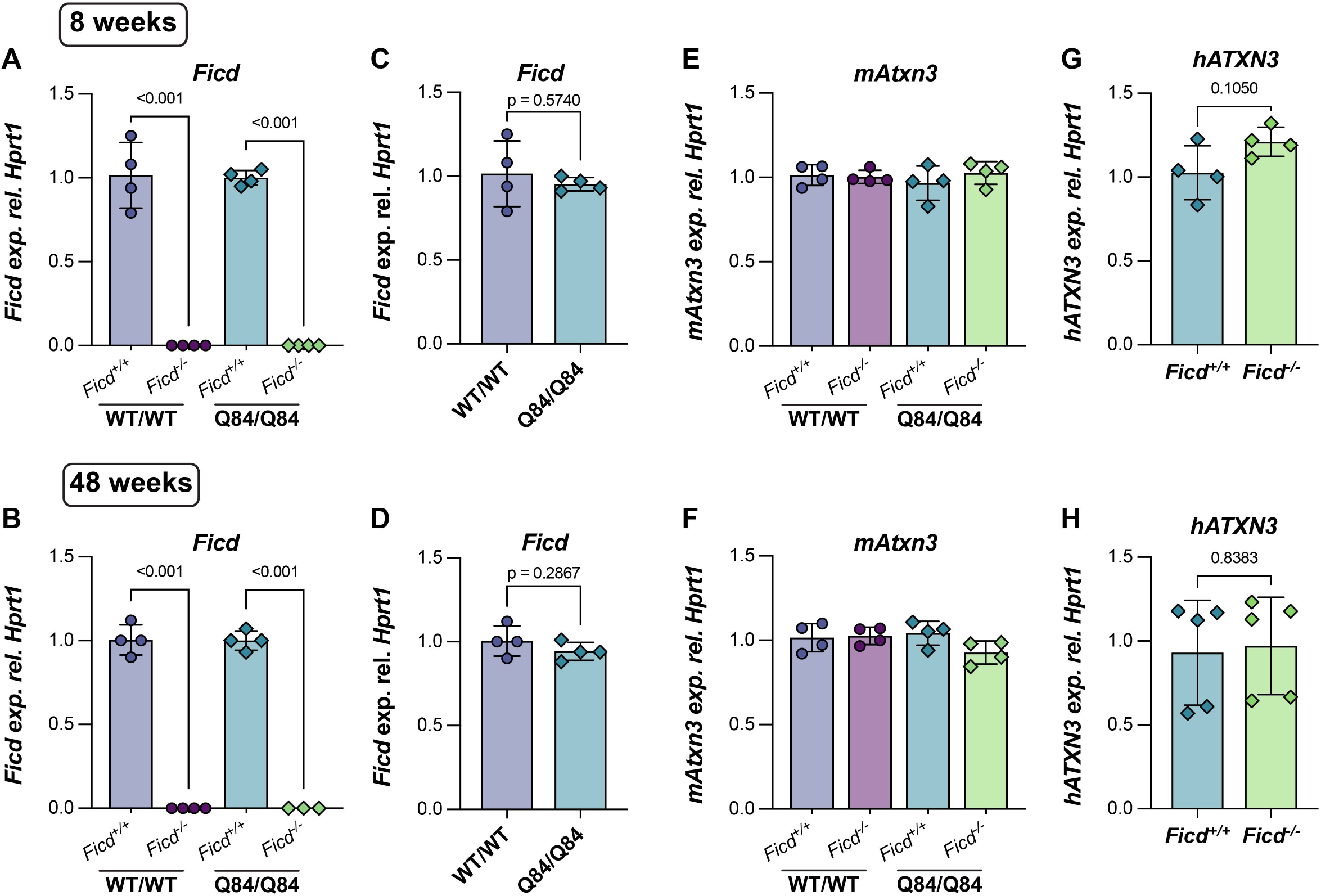
(A-B) *Ficd* transcript levels in the brainstem of 8 (A) and 48 week-old (B) *Ficd*^-/-^ mice normalized to WT/WT and Q84/Q84; *Ficd*^+/+^ littermate controls, respectively. (C-D) *Ficd* expression in Q84/Q84; *Ficd*^+/+^ mouse brainstem at 8 (C) and 48 weeks (D) normalized to WT/WT; *Ficd*^+/+^ littermates. (E-F) Brainstem transcript levels of endogenous mouse *Atxn3 (mAtxn3)* at 8 (E) and 48 weeks (F) in all four experimental groups. (G-H) Expression levels of the mutant human *ATXN3* transgene (*hATXN3*) in the brainstem of Q84/Q84; *Ficd*^+/+^ and Q84/Q84; *Ficd*^-/-^ mice at 8 (G) and 48 (H) weeks of age. A minimum of *n* = 3 animals per experimental group are displayed in each plot. Data are presented as mean ± SD. For (A-B, E-F), statistical significance was determined by one-way ANOVA followed by Tukey’s post-hoc multiple comparisons testing. For (C-D, G-H), Welch’s t-test was used to compute statistical significance.

**Fig. S2.**
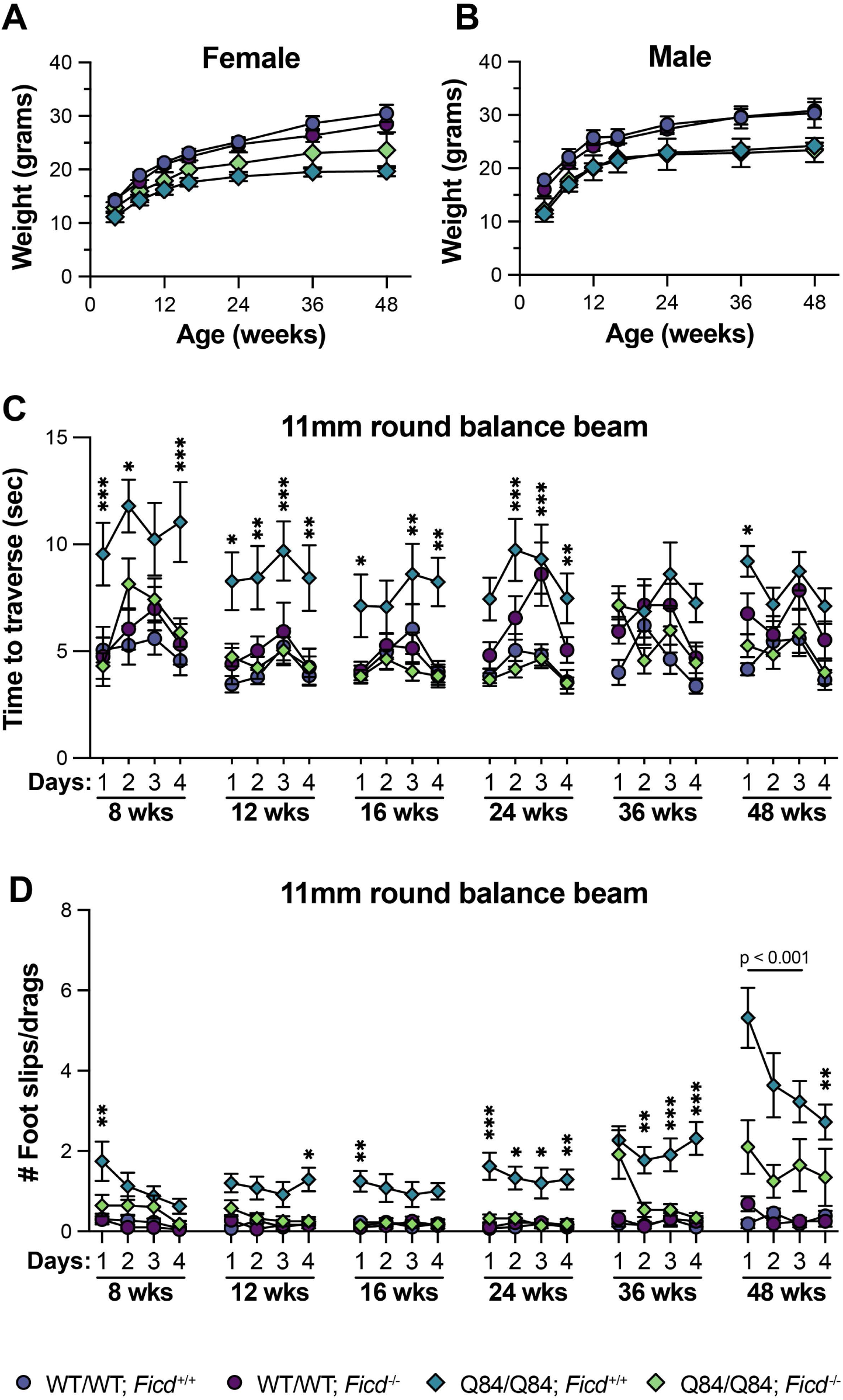
(A-B) Graphs showing the weight in grams of (A) female and (B) male mice from each experimental group recorded at weaning (4 weeks) and at each behavioral testing time-point through 48 weeks. (C) Average time taken to traverse the 11mm round balance beam on each of 4 consecutive trial days for all time-points tested. (D) Average number of hindlimb foot slips and/or drags during the 11mm round balance beam traversal task. For (A-D), data are shown as mean ± SEM. For (C-D), statistical significance was determined by two-way ANOVA followed by Tukey’s post-hoc multiple comparisons testing. P-values reflect comparisons between Q84/Q84; *Ficd*^+/+^ and Q84/Q84; *Ficd*^-/-^ groups. *p<0.05; **p<0.01; ***p<0.001. A minimum of 4 male and 4 female mice per experimental group were used in all behavioral studies.

**Fig. S3.**
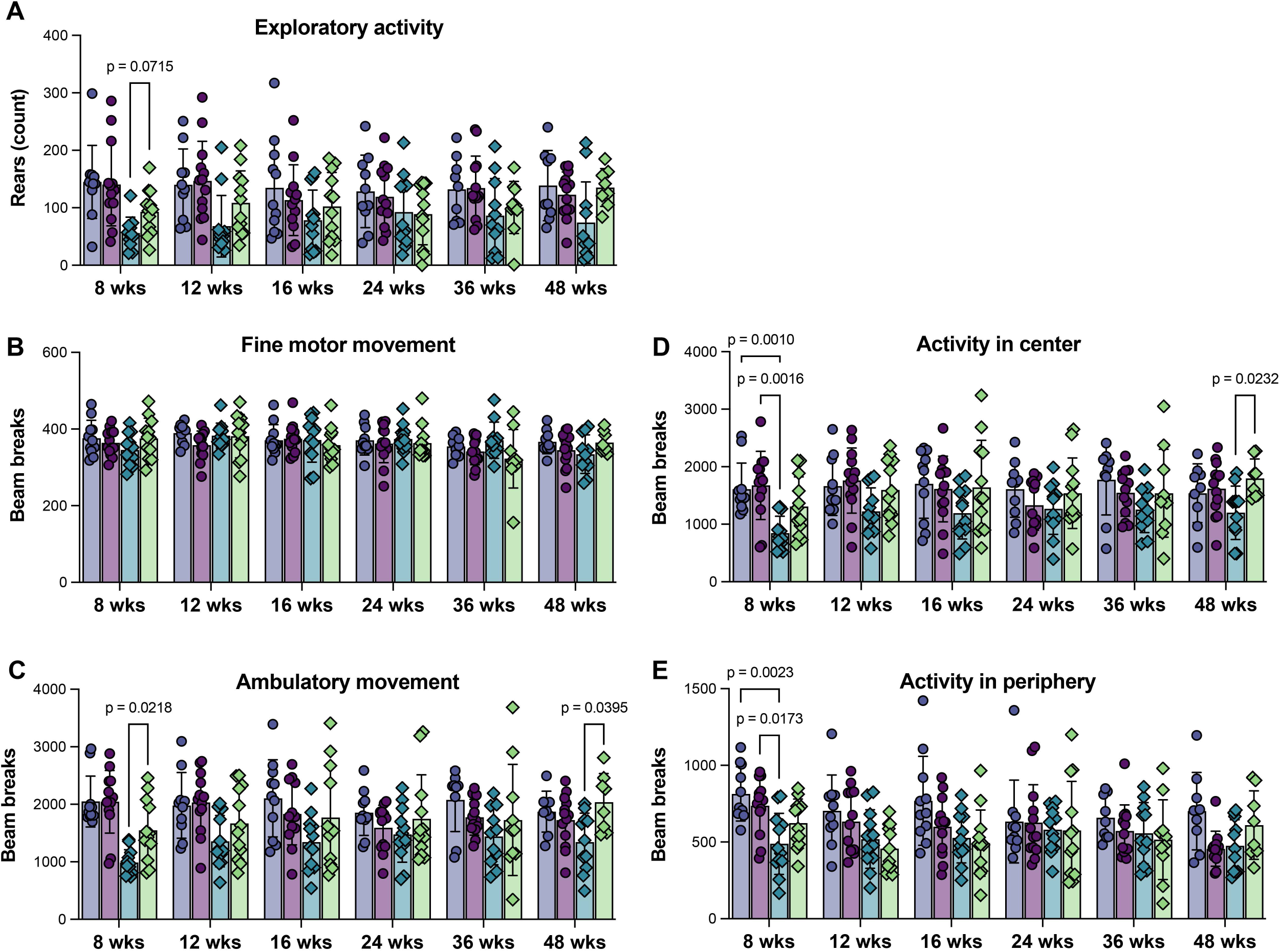
(A) Mouse exploratory activity determined by the number of rearing events recorded during the 30 minute open field session. (B-C) Fine motor movement (B) and ambulatory movement (C) in the open field chamber assessed by the number of photobeam breaks. (D-E) Amount of activity recorded in the center (D) or periphery (E) of the open field chamber. Data are displayed as mean ± SD. Statistical significance was determined using a mixed-effects analysis followed by Tukey’s post-hoc multiple comparisons testing. A minimum of 4 male and 4 female mice per experimental group were used in all behavioral studies.

**Fig. S4.**
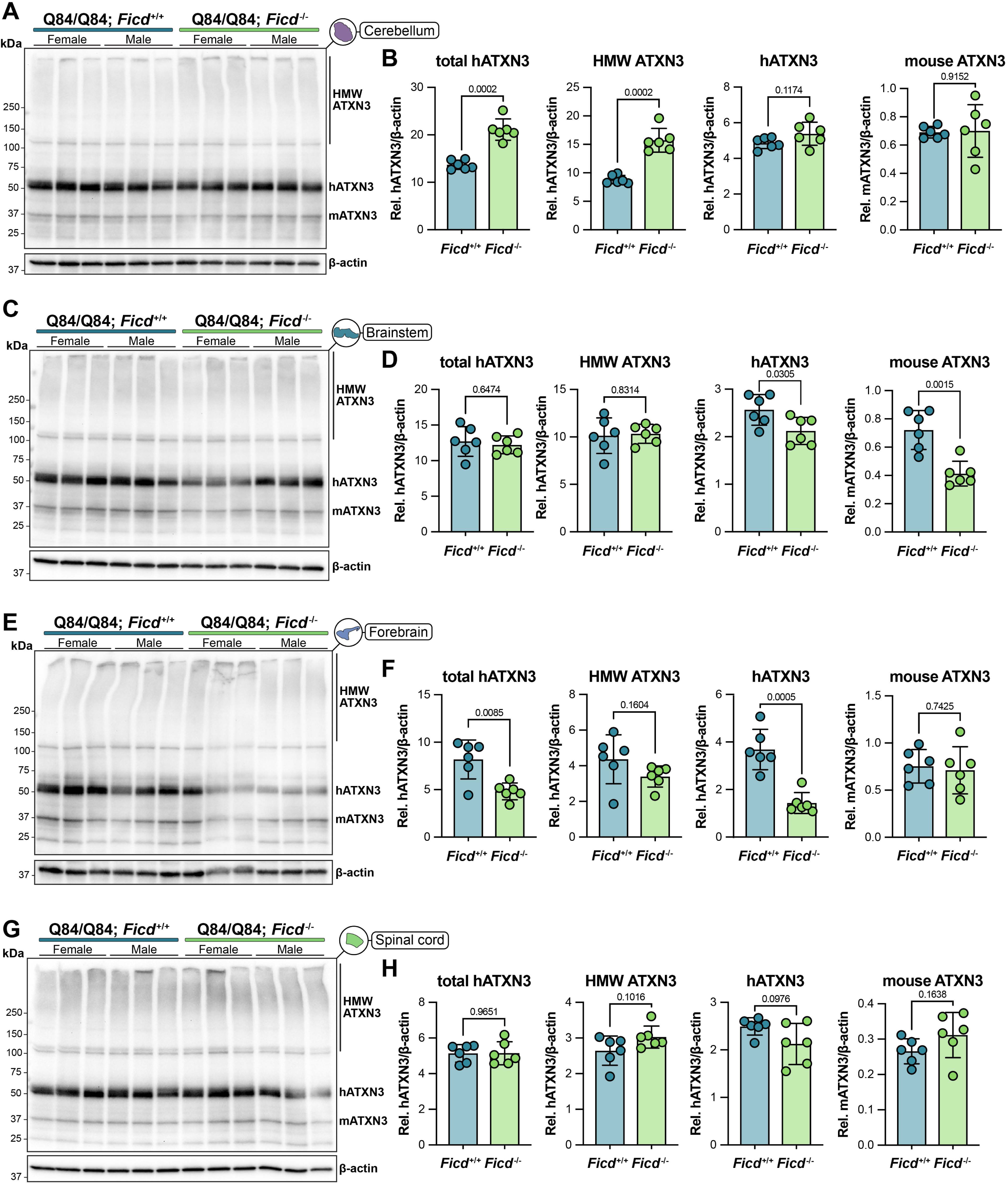
(A-B) Representative western blot (A) of ATXN3 levels in the cerebellum of 8 week-old mice and (B) quantification of total (HMW + monomeric), high molecular weight (HMW), and monomeric mutant human ATXN3 (hATXN3) as well as endogenous mouse ATXN3 (mATXN3). (C-D) Representative western blot and quantification of ATXN3 levels in the brainstem of 8 week-old mice. (E-F) Representative western blot and quantification of ATXN3 levels in the forebrain of 8 week-old mice. (G-H) Representative western blot and quantification of ATXN3 levels in the spinal cord of 8 week-old mice. Data are mean ± SD. Samples from a total of 3 male and 3 female mice (*n* = 6) per genotype were used in all experiments. Statistical significance was determined using unpaired parametric *t* tests with exact p-values reported on each graph.

**Fig. S5.**
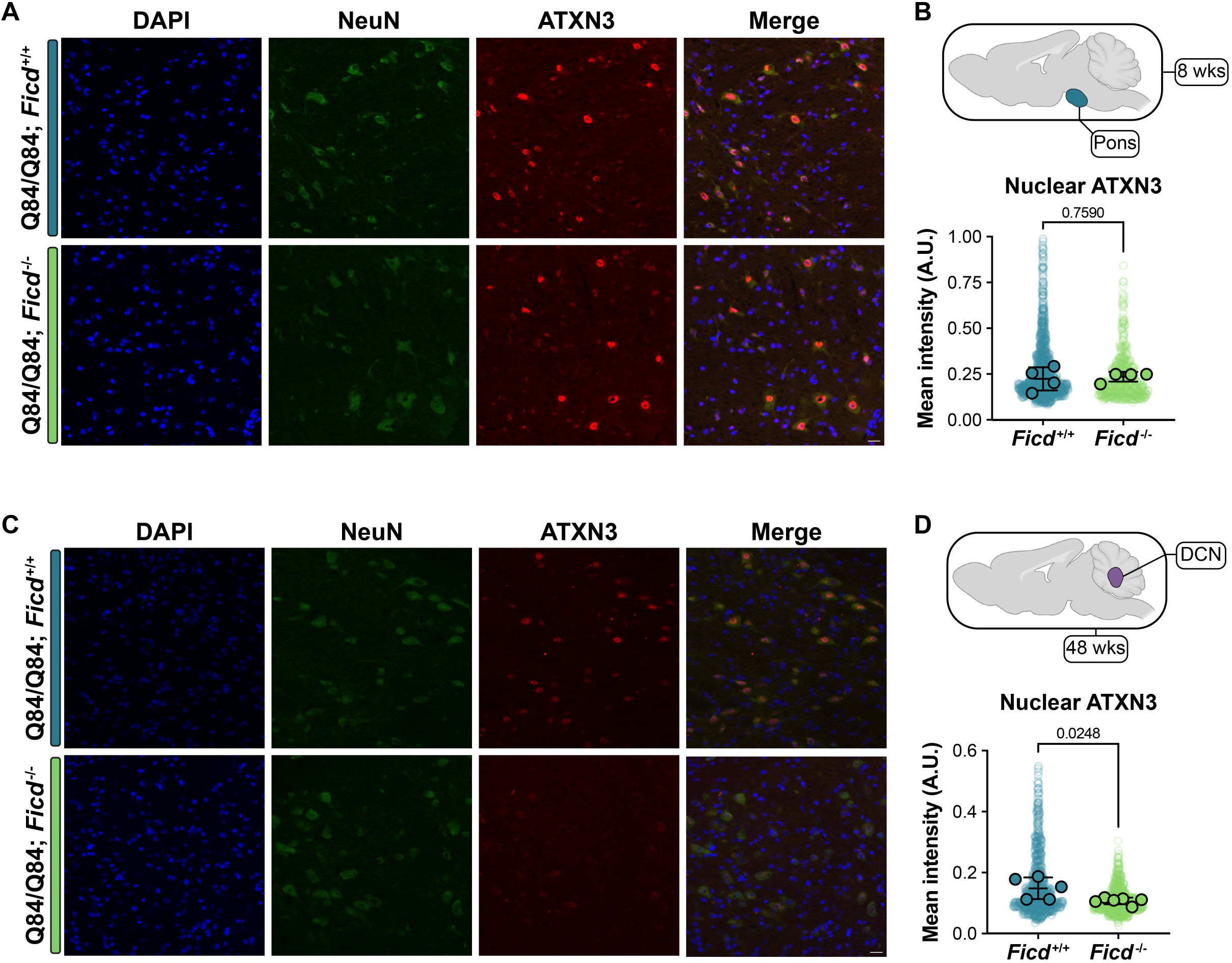
(A) Representative sagittal immunofluorescent images of DAPI (blue), NeuN (green), and ATXN3 (red) expression in the pons of 8 week-old mice. Scale bar = 20 μm. (B) Quantification of nuclear ATXN3 levels in NeuN+ cells (*n* = 4 images per mouse, 2 male and 2 female mice per genotype). (C) Representative sagittal immunofluorescent images of DAPI (blue), NeuN (green), and ATXN3 (red) expression in the deep cerebellar nuclei (DCN) of 48 week-old mice. (D) Quantification of nuclear ATXN3 levels in NeuN+ cells (*n* = 6 images per mouse, at least 5 mice per genotype). For (B, D) translucent data points represent individual cells, while opaque data points reflect the average intensity for each animal. Statistical significance was computed by performing unpaired parametric *t* tests on per-mouse averages, with exact p-values displayed on graphs.

**Fig. S6.**
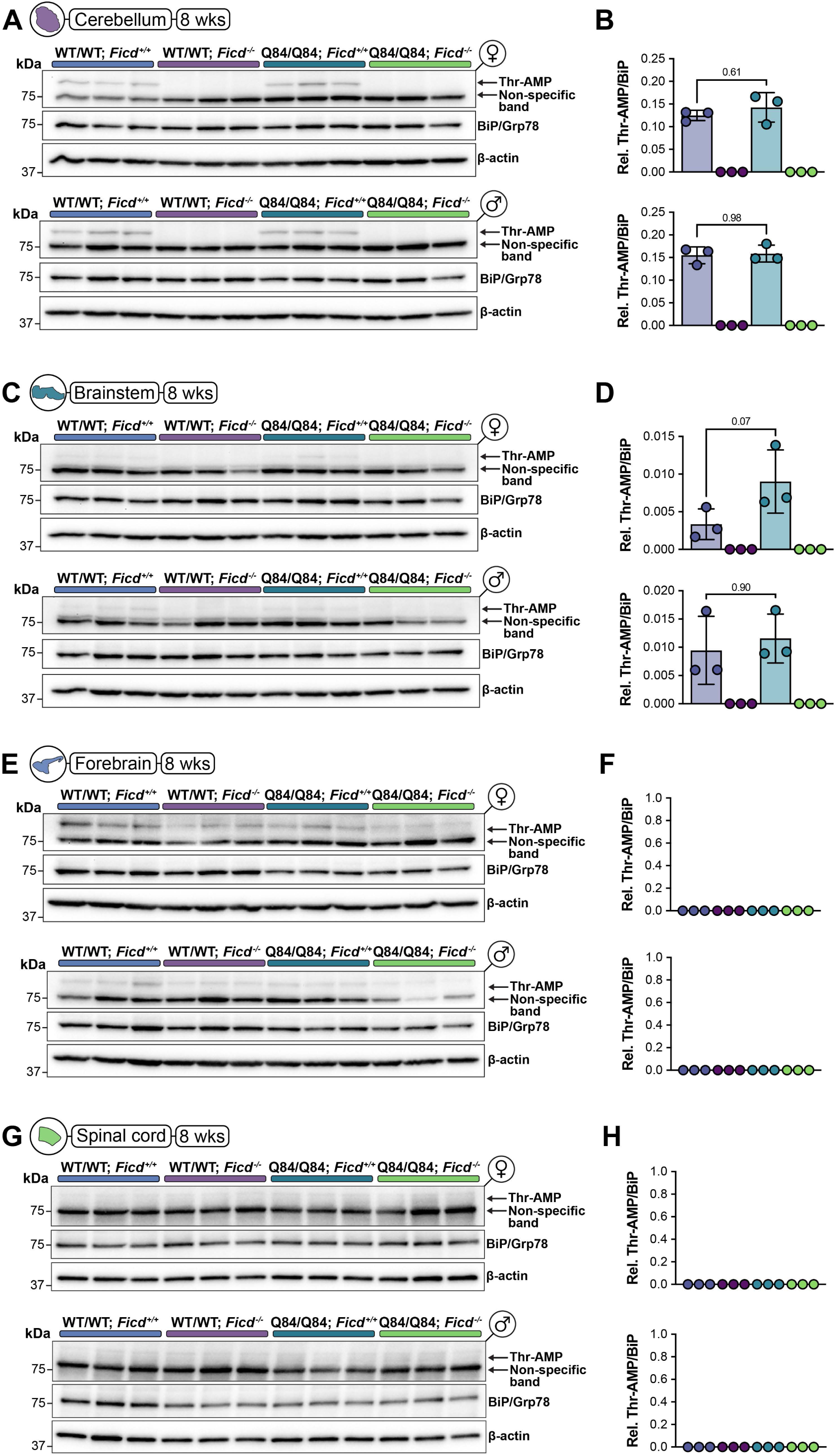
(A-B) Western blot (A) showing levels of AMPylated threonine (Thr-AMP), BiP, and β-actin in the cerebellum of 8 week-old female (top) and male (bottom) mice from all 4 experimental groups and (B) quantification of AMPylated BiP levels normalized to total BiP. (C-D) Western blot (C) and (D) quantification of AMPylated BiP levels in the brainstem of female (top) and male (bottom) mice. (E-F) Western blot (E) and (F) quantification of AMPylated BiP levels in the forebrain of female (top) and male (bottom) mice. (G-H) Western blot (G) and (H) quantification of AMPylated BiP levels in the cervical spinal cord of female (top) and male (bottom) mice. For each genotype group, *n* = 6 mice (3 male, 3 female) were used in all experiments. Data are shown as mean ± SD. Statistical significance was determined by one-way ANOVA with Tukey’s post-hoc multiple comparisons testing. Exact p-values are displayed on graphs.

**Fig. S7.**
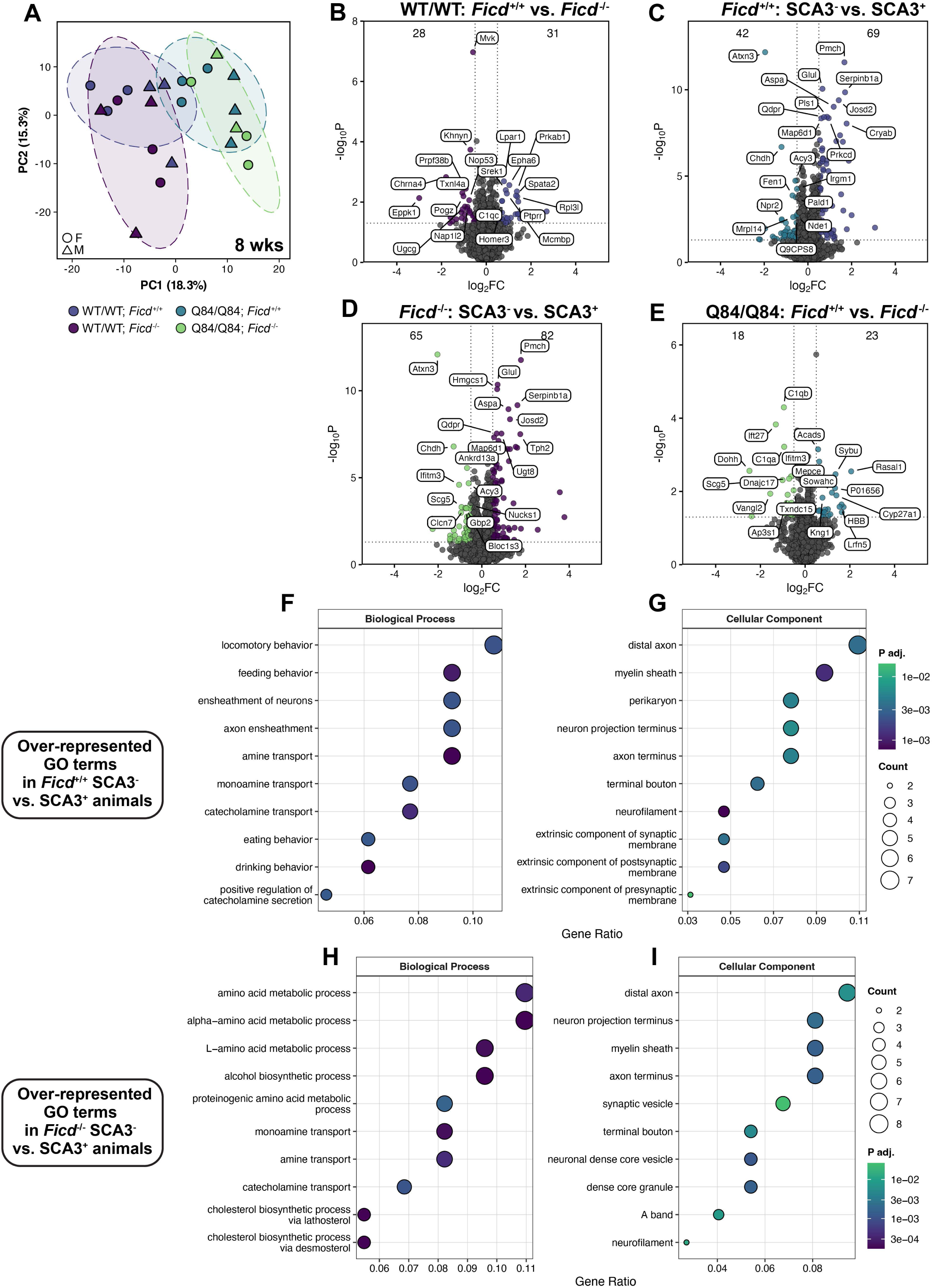
(A) Principal component analysis (PCA) plot showing clustering of 8 week-old samples by genotype. Each data point reflects one mouse. Females are circles, males are triangles. (B) Volcano plot depicting differentially-expressed (DE) proteins in SCA3-negative (WT/WT) *Ficd*^+/+^ vs. *Ficd*^-/-^ cohorts. (C) Volcano plot comparing SCA3-negative vs. SCA3-positive (Q84/Q84) *Ficd*^+/+^ groups. (D) Volcano plot comparing SCA3-negative vs. SCA3-positive (Q84/Q84) *Ficd*^-/-^ groups. (E) Volcano plot comparing Q84/Q84; *Ficd*^+/+^ vs. *Ficd*^-/-^ proteomes. For (B-E), dashed lines reflect statistical cut-offs (log_2_FC > |0.5|, p < 0.05). Top DE proteins for each group are labeled, while numbers at the top of each graph indicate the total number of DE proteins. (F-G) Dot plots showing the top biological process (F) and cellular compartment (G) gene ontology (GO) terms over-represented in SCA3-negative vs. SCA3-positive *Ficd*^+/+^ animals. (H-I) Dot plots showing the top biological process (H) and cellular compartment (I) gene ontology (GO) terms over-represented in SCA3-negative vs. SCA3-positive *Ficd*^+/+^ animals. For (F-I), dot size reflects the number of proteins associated with the GO term and dots are colored according to adjusted p-value (*P_adj_*).

**Fig. S8.**
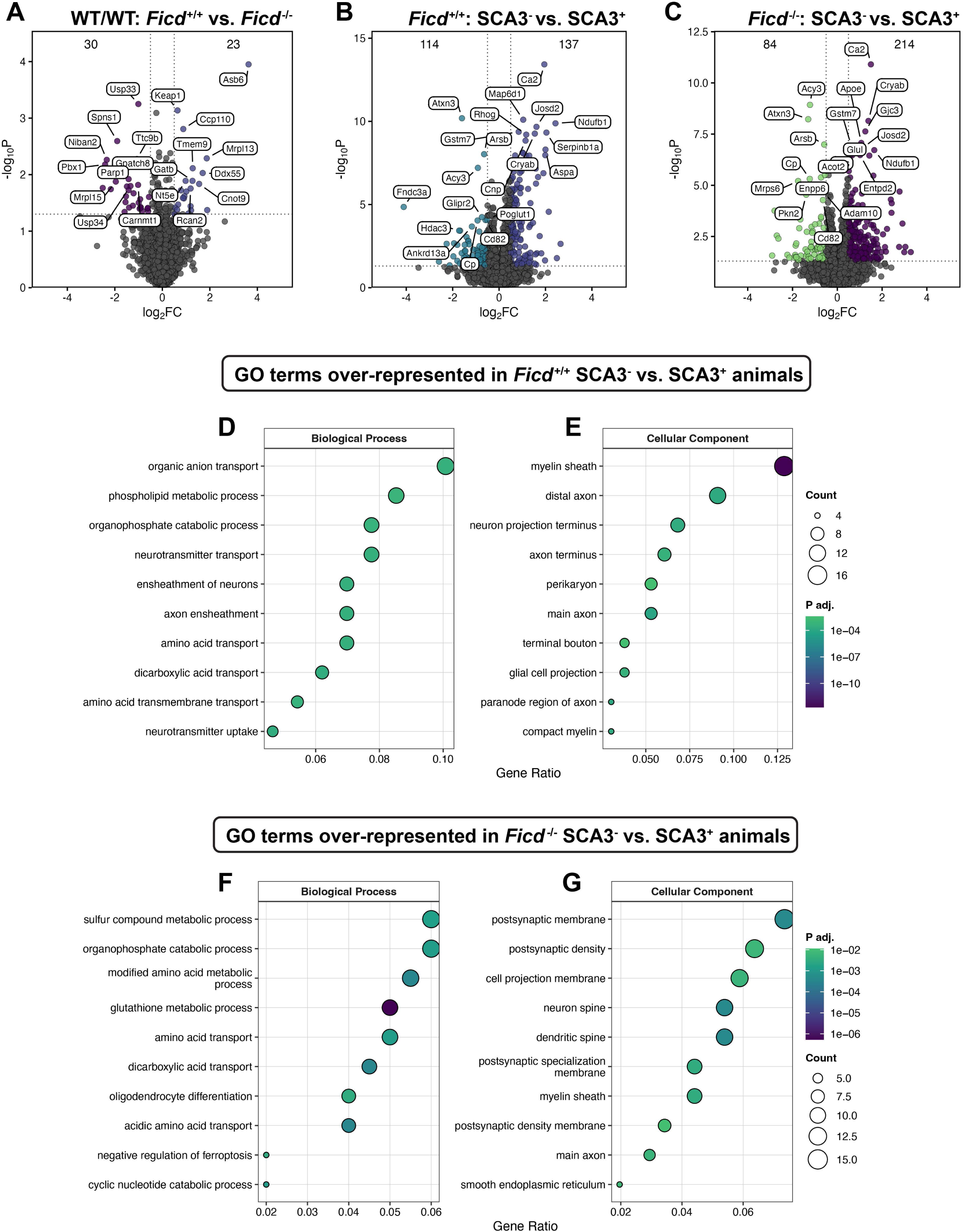
(A) Volcano plot depicting differentially-expressed (DE) proteins in SCA3-negative (WT/WT) *Ficd*^+/+^ vs. *Ficd*^-/-^ cohorts at 48 weeks. (B) Volcano plot comparing SCA3-negative vs. SCA3-positive (Q84/Q84) *Ficd*^+/+^ groups. (C) Volcano plot comparing SCA3-negative vs. SCA3-positive (Q84/Q84) *Ficd*^-/-^ groups. For (A-C), dashed lines reflect statistical cut-offs (log_2_FC > |0.5|, p < 0.05). Top DE proteins for each group are labeled, while numbers at the top of each graph indicate the total number of DE proteins in that group. (D-E) Dot plots showing the top biological process (D) and cellular compartment (E) gene ontology (GO) terms over-represented in SCA3-negative vs. SCA3-positive *Ficd*^+/+^ proteomes. (F-G) Dot plots showing the top biological process (F) and cellular compartment (G) gene ontology (GO) terms over-represented in SCA3-negative vs. SCA3-positive *Ficd*^+/+^ animals. For (D-G), dot size reflects the number of proteins associated with the GO term and dots are colored according to adjusted p-value (*P_adj_*).

**Fig. S9.**
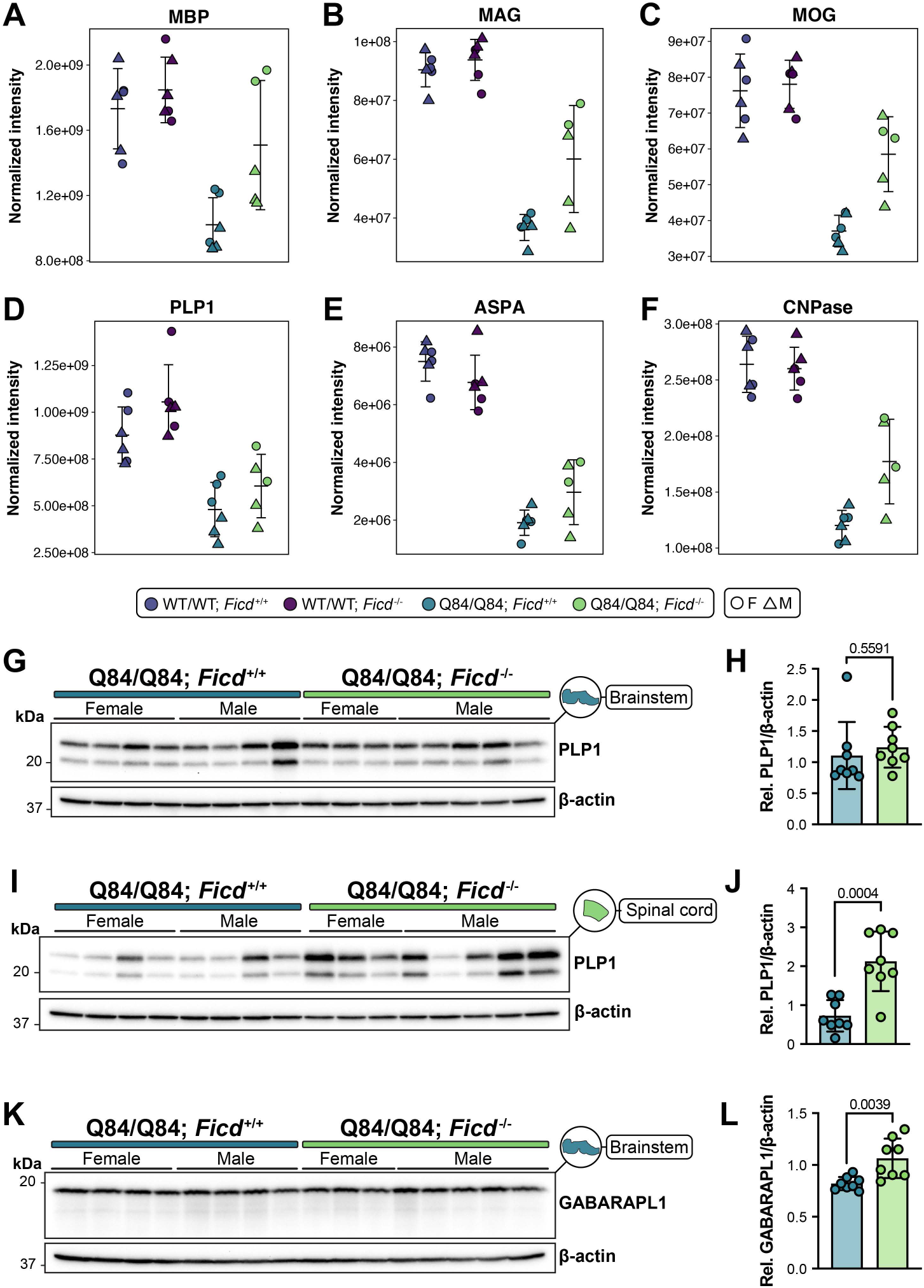
(A-F) Normalized intensity values from DIA-MS data of (A) MBP, (B) MAG, (C) MOG, (D) PLP1, (E) ASPA, and (F) CNPase from all four experimental genotypes. Data are mean ± SD. One data point represents one mouse. Sex is indicated by symbol type. (G) Western blot and (H) quantification of proteolipid protein 1 (PLP1) levels in the brainstem of 48 week-old mice. (I) Western blot and (J) quantification of PLP1 levels in the spinal cord of 48 week-old mice. The beta-actin loading control in (I) is the same loading control shown in Fig. 5G. The same membrane was stripped and re-probed for each target. The loading control is intentionally shown in both figures for clarity. (K) Western blot and (L) quantification of GABARAPL1 levels in the brainstem of 48 week-old mice. Data are mean ± SD. For all experiments, *Ficd*^+/+^ *n* = 4 female, 4 male mice, and *Ficd*^-/-^ *n* = 3 female, 5 male mice. Statistical significance was determined by unpaired, parametric *t* tests. Exact p-values displayed on graphs

**Fig. S10.**
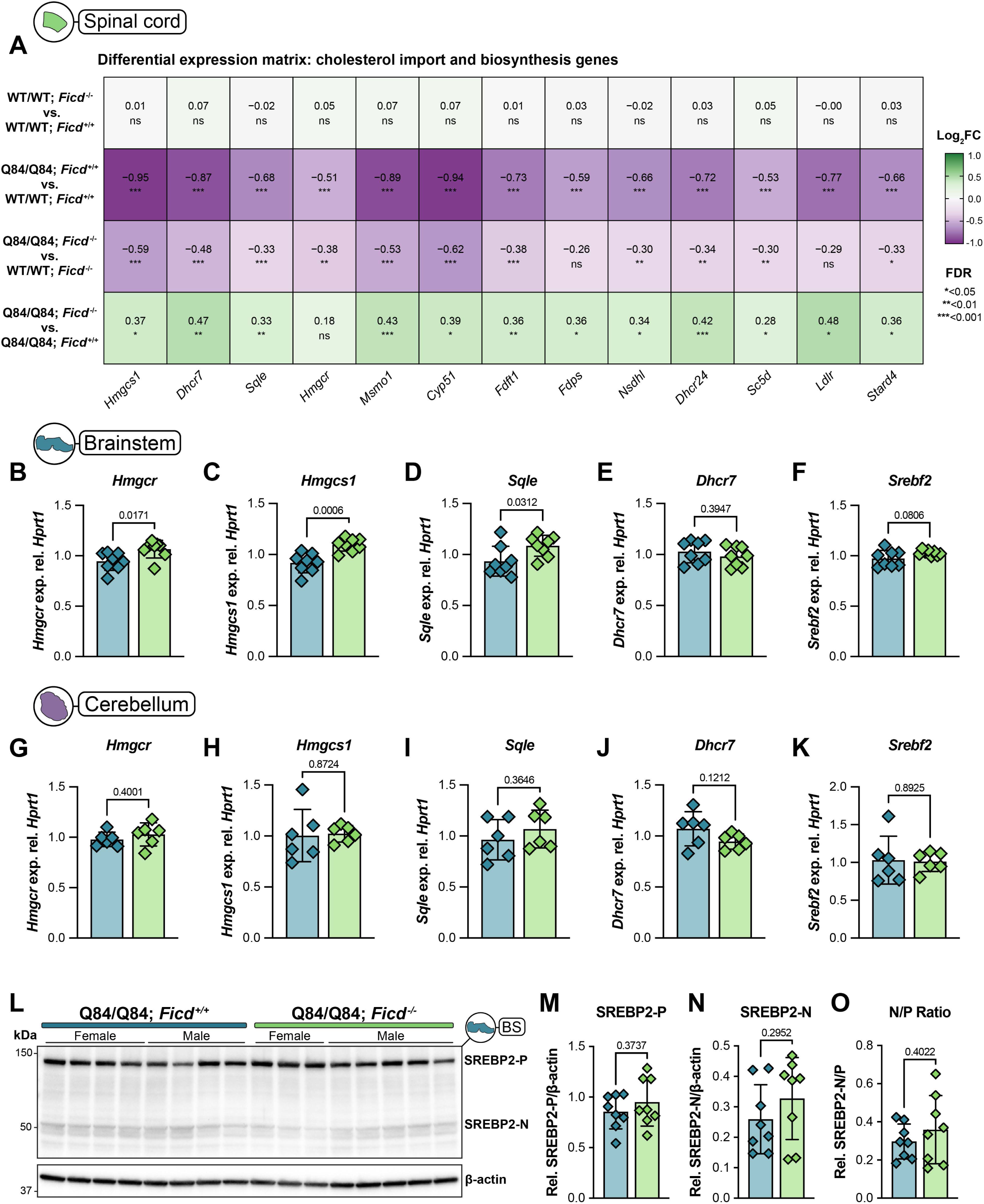
(A) Heatmap showing directional log_2_ fold-change values in the spinal cord for cholesterol biosynthesis and import genes across experimental contrasts. Statistical significance determined by edgeR likelihood ratio tests (*FDR<0.05, **FDR<0.01, ***FDR<0.001). (B-F) Quantitative PCR (qPCR) assessing expression levels of key cholesterol synthesis genes (B) *Hmgcr*, (C) *Hmgcs1*, (D) *Sqle*, (E) *Dhcr7*, and (F) *Srebf2* in the brainstem of 48 week-old mice (*n* = 6 mice per genotype, split evenly by sex). Data are mean ± SD. (G-K) Expression levels of (G) *Hmgcr*, (H) *Hmgcs1*, (I) *Sqle*, (J) *Dhcr7*, and (K) *Srebf2* in the cerebellum of 48 week-old mice (*n* = 6 mice per genotype, split evenly by sex). Data are mean ± SD. (L) Western blot depicting levels of precursor (-P) and nuclear (-N) SREBP2 in the brainstem (*Ficd*^+/+^ *n* = 4 male, 4 female mice; *Ficd*^-/-^ *n* = 3 female, 5 male mice). (M-O) Quantification of (M) precursor (SREBP2-P), (N) nuclear (SREBP2-N), and (O) ratio of nuclear to precursor SREBP2. Data are mean ± SD. For (B-K, M-O) statistical significance was determined by unpaired, parametric *t* tests. Exact p-values displayed on graphs.

**Table S1.**
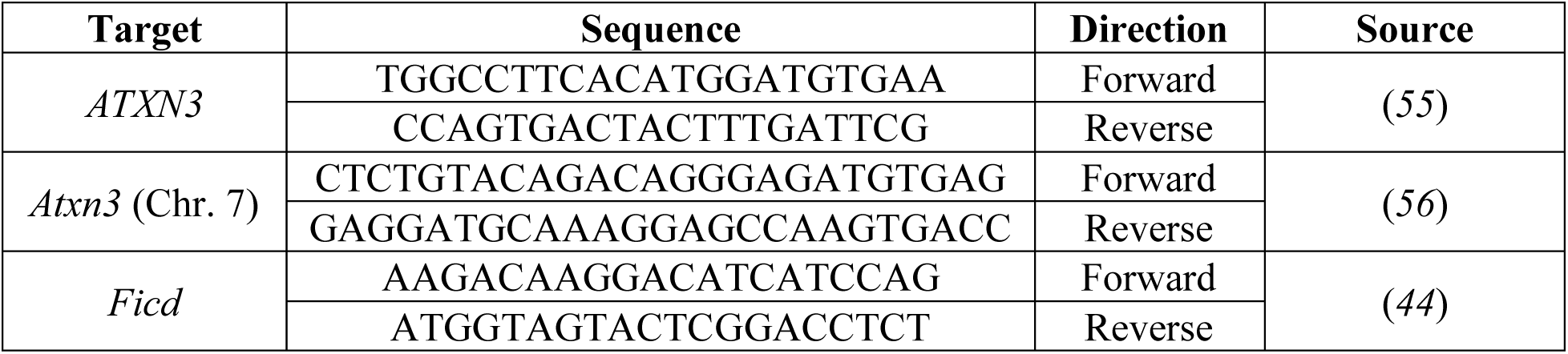
PCR primers used for YAC-Q84 x *Ficd^-/-^*mouse genotyping.

**Table S2.** Mouse IDs and CAG repeat sizes of 8 week-old mice.

| Genotype | Gender | Mouse ID | CAG repeat size |
| --- | --- | --- | --- |
| WT/WT<br>*Ficd*<sup>+/+</sup> | F | 38.3.11 | N/A |
|  |  | 38.4.19 |  |
|  |  | 46.1.4 |  |
|  | M | 33.1.4 |  |
|  |  | 38.2.8 |  |
|  |  | 37.5.17 |  |
|  |  | 37.6.22 |  |
| WT/WT<br>*Ficd*<sup>-/-</sup> | F | 32.3.13 | N/A |
|  |  | 35.1.1 |  |
|  |  | 36.1.2 |  |
|  |  | 33.2.9 |  |
|  | M | 32.1.5 |  |
|  |  | 34.2.10 |  |
|  |  | 35.2.6 |  |
|  |  | 35.2.9 |  |
| Q84/Q84<br>*Ficd*<sup>+/+</sup> | F | 32.3.10 | 78 |
|  |  | 38.1.2 | 74 |
|  |  | 33.2.6 | 75 |
|  |  | 37.5.13 | 75 |
|  | M | 32.1.4 | 78 |
|  |  | 38.2.6 | 73 |
|  |  | 38.3.15 | 75 |
|  |  | 38.3.16 | 72 |
|  |  | 46.1.6 | 79 |
| Q84/Q84<br>*Ficd*<sup>-/-</sup> | F | 32.1.1 | 80 |
|  |  | 28.5.20 | 77 |
|  |  | 34.2.5 | 75 |
|  |  | 38.2.5 | 72 |
|  | M | 33.3.8 | 79 |
|  |  | 32.5.24 | 76 |
|  |  | 38.5.25 | 73 |

**Table S3.**
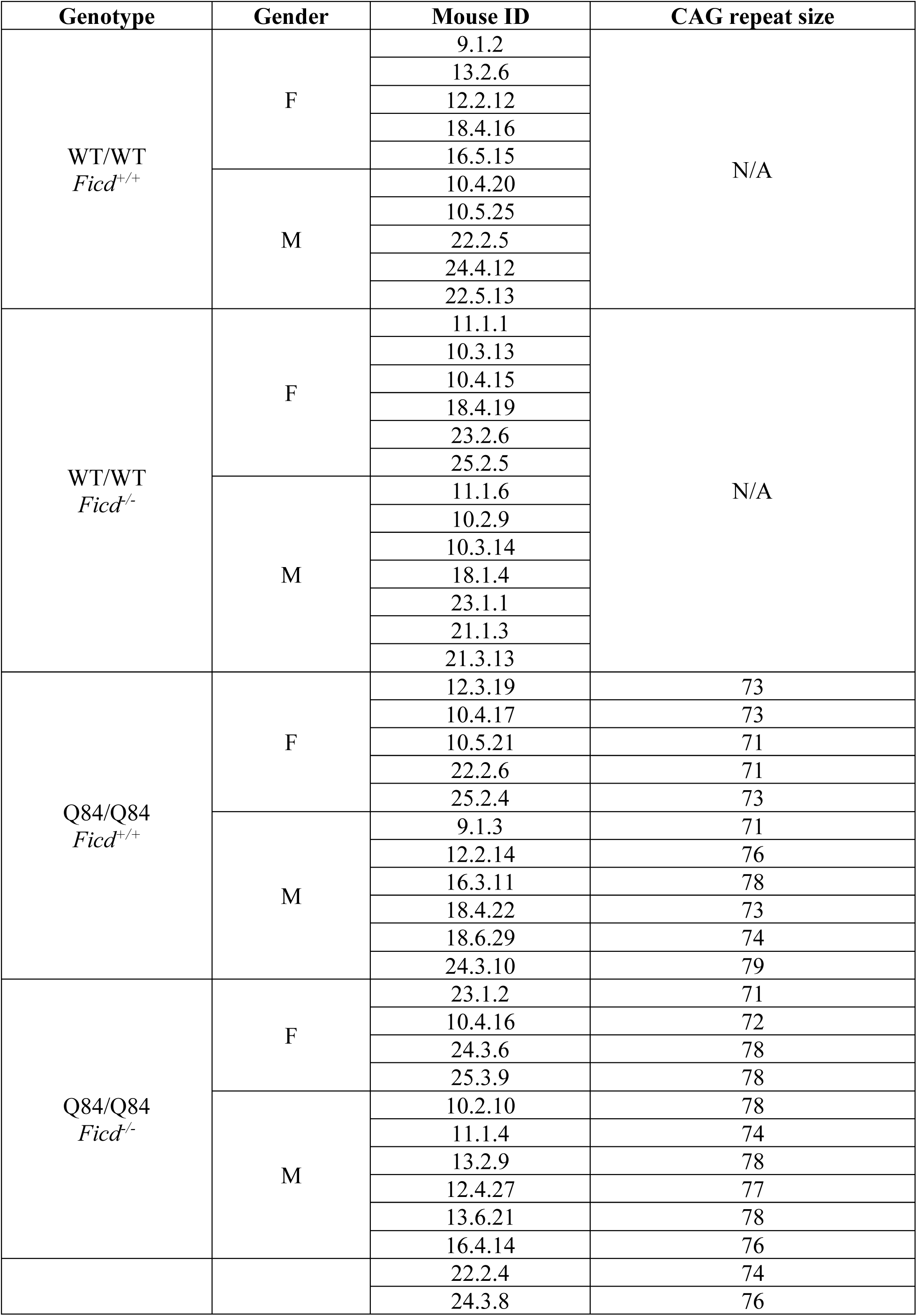
Mouse IDs and CAG repeat sizes of 48 week-old mice.

**Table S4.** Primary antibodies used for western blots in this study.

| **Target** | **Host** | **Dilution** | **Manufacturer** | **Cat #** |
| --- | --- | --- | --- | --- |
| MJD (ATXN3) | Rabbit | 1:30000 | (107) | N/A |
| Thr-AMP | Mouse | 1:2000 | Biointron / (58) | N/A |
| Vinculin | Rabbit | 1:2000 | Cell Signaling Technology | 23214 |
| Beta-actin | Mouse | 1:5000 | Cell Signaling Technology | 3700 |
| GAPDH | Mouse | 1:5000 | Proteintech | 60004-1-Ig |
| BIP/GRP78 | Mouse | 1:2000 | Proteintech | 66574-1-Ig |
| MBP | Mouse | 1:1000 | Santa Cruz Biotechnology | sc-66064 |
| PLP1 | Rabbit | 1:1000 | Cell Signaling Technology | 28702 |
| GABARAPL1 | Rabbit | 1:1000 | Cell Signaling Technology | 26632 |
| SREBP2 | Rabbit | 1:500 | Invitrogen | PA1-338 |
| Aspartoacylase | Rabbit | 1:1000 | Cell Signaling Technology | 25959 |
| MAG | Rabbit | 1:1000 | Cell Signaling Technology | 9043 |

**Table S5.** Window widths for DIA-MS data acquisition.

| **m/z** | **Window width (m/z)** |
| --- | --- |
| 377 | 54 |
| 419 | 32 |
| 448 | 28 |
| 473.5 | 25 |
| 497.5 | 25 |
| 520.5 | 23 |
| 542.5 | 23 |
| 564.5 | 23 |
| 587 | 24 |
| 610.5 | 25 |
| 635 | 23 |
| 660 | 23 |
| 685.5 | 27 |
| 712.5 | 29 |
| 741 | 30 |
| 771 | 32 |
| 803.5 | 35 |
| 838.5 | 37 |
| 877 | 42 |
| 921 | 48 |
| 972 | 52 |
| 1034.5 | 71 |
| 113.5 | 129 |
| 1423.5 | 453 |

**Table S6.**
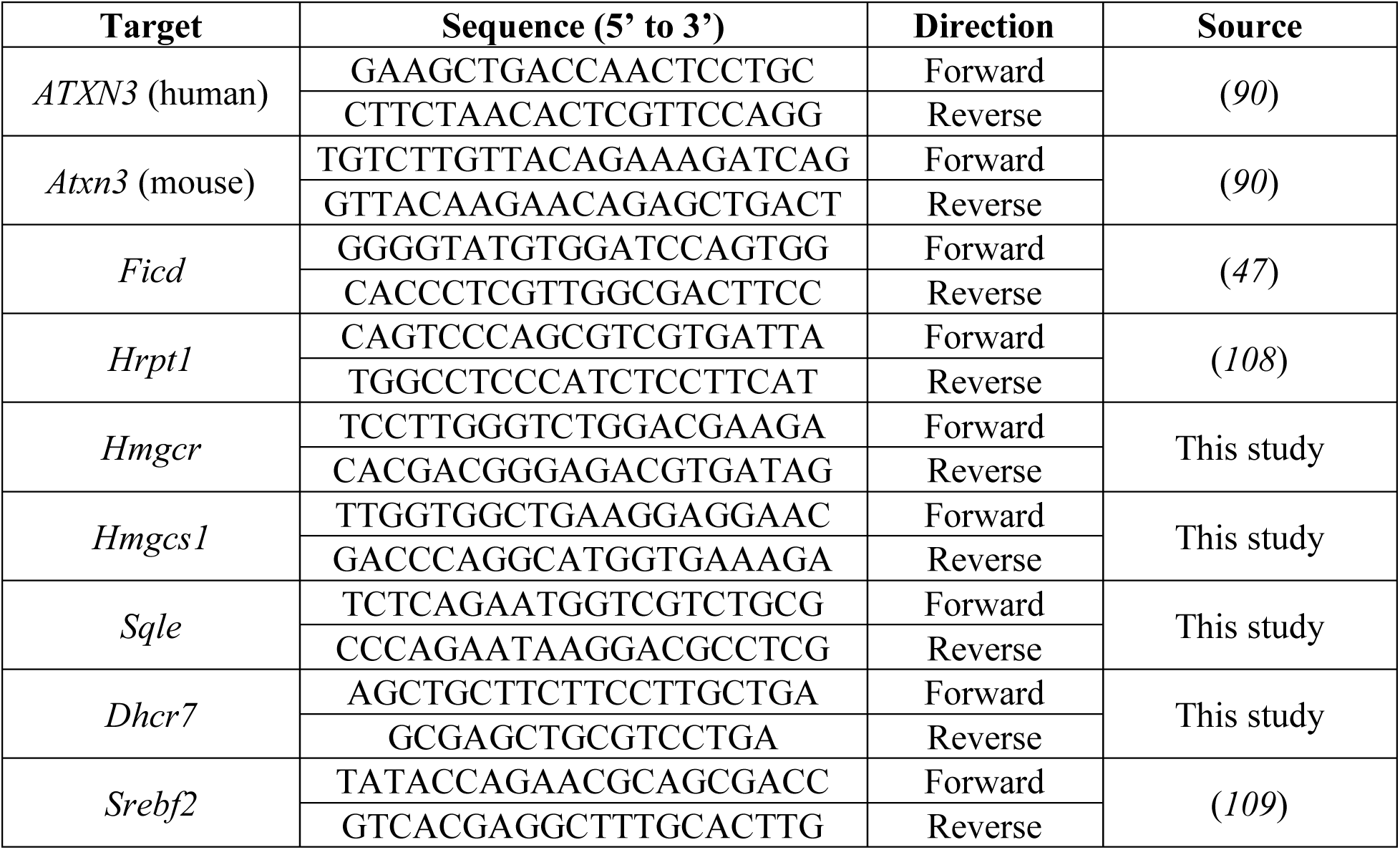
RT-qPCR primers used in this study.

## REFERENCES

1. A. P. Lieberman, V. G. Shakkottai, R. L. Albin, Polyglutamine Repeats in Neurodegenerative Diseases. Annu Rev Pathol 14, 1–27 (2019).

2. H. S. McLoughlin, L. R. Moore, H. L. Paulson, Pathogenesis of SCA3 and implications for other polyglutamine diseases. Neurobiology of Disease 134, 104635 (2020).

3. H. L. Paulson, V. G. Shakkottai, H. B. Clark, H. T. Orr, Polyglutamine spinocerebellar ataxias - from genes to potential treatments. Nat Rev Neurosci 18, 613–626 (2017).

4. H. Paulson, Machado-Joseph Disease/Spinocerebellar Ataxia Type 3. Handb Clin Neurol 103, 437–449 (2012).

5. W. Scherzed, E. R. Brunt, H. Heinsen, R. A. de Vos, K. Seidel, K. Bürk, L. Schöls, G. Auburger, D. Del Turco, T. Deller, H. W. Korf, W. F. den Dunnen, U. Rüb, Pathoanatomy of cerebellar degeneration in spinocerebellar ataxia type 2 (SCA2) and type 3 (SCA3). Cerebellum 11, 749–760 (2012).

6. U. Rüb, E. R. Brunt, T. Deller, New insights into the pathoanatomy of spinocerebellar ataxia type 3 (Machado-Joseph disease). Curr Opin Neurol 21, 111–116 (2008).

7. M.-L. Monin, S. Tezenas du Montcel, C. Marelli, C. Cazeneuve, P. Charles, C. Tallaksen, S. Forlani, G. Stevanin, A. Brice, A. Durr, Survival and severity in dominant cerebellar ataxias. Ann Clin Transl Neurol 2, 202–207 (2015).

8. C. A. Matos, L. P. de Almeida, C. Nóbrega, Machado–Joseph disease/spinocerebellar ataxia type 3: lessons from disease pathogenesis and clues into therapy. Journal of Neurochemistry 148, 8–28 (2019).

9. F. E. Reyes-Turcu, K. H. Ventii, K. D. Wilkinson, Regulation and Cellular Roles of Ubiquitin-Specific Deubiquitinating Enzymes. Annual Review of Biochemistry 78, 363–397 (2009).

10. K. M. Scaglione, E. Zavodszky, S. V. Todi, S. Patury, P. Xu, E. Rodríguez-Lebrón, S. Fischer, J. Konen, A. Djarmati, J. Peng, J. E. Gestwicki, H. L. Paulson, Ube2w and Ataxin-3 Coordinately Regulate the Ubiquitin Ligase CHIP. Molecular Cell 43, 599–612 (2011).

11. H. L. Paulson, M. K. Perez, Y. Trottier, J. Q. Trojanowski, S. H. Subramony, S. S. Das, P. Vig, J. L. Mandel, K. H. Fischbeck, R. N. Pittman, Intranuclear inclusions of expanded polyglutamine protein in spinocerebellar ataxia type 3. Neuron 19, 333–344 (1997).

12. L. S. Havel, S. Li, X.-J. Li, Nuclear accumulation of polyglutamine disease proteins and neuropathology. Mol Brain 2, 21 (2009).

13. L. J. A. Toonen, M. Overzier, M. M. Evers, L. G. Leon, S. A. J. van der Zeeuw, H. Mei, S. M. Kielbasa, J. J. Goeman, K. M. Hettne, O. Th. Magnusson, M. Poirel, A. Seyer, P. A. C. ‘t Hoen, W. M. C. van Roon-Mom, Transcriptional profiling and biomarker identification reveal tissue specific effects of expanded ataxin-3 in a spinocerebellar ataxia type 3 mouse model. Mol Neurodegener 13, 31 (2018).

14. B. Ramani, B. Panwar, L. R. Moore, B. Wang, R. Huang, Y. Guan, H. L. Paulson, Comparison of spinocerebellar ataxia type 3 mouse models identifies early gain-of-function, cell-autonomous transcriptional changes in oligodendrocytes. Hum Mol Genet 26, 3362–3374 (2017).

15. K. H. Schuster, A. J. Zalon, H. Zhang, D. M. DiFranco, N. R. Stec, Z. Haque, K. G. Blumenstein, A. M. Pierce, Y. Guan, H. L. Paulson, H. S. McLoughlin, Impaired Oligodendrocyte Maturation Is an Early Feature in SCA3 Disease Pathogenesis. J. Neurosci. 42, 1604–1617 (2022).

16. E. Haas, R. D. Incebacak, T. Hentrich, C. Huridou, T. Schmidt, N. Casadei, Y. Maringer, C. Bahl, F. Zimmermann, J. D. Mills, E. Aronica, O. Riess, J. M. Schulze-Hentrich, J. Hübener-Schmid, A Novel SCA3 Knock-in Mouse Model Mimics the Human SCA3 Disease Phenotype Including Neuropathological, Behavioral, and Transcriptional Abnormalities Especially in Oligodendrocytes. Mol Neurobiol 59, 495–522 (2022).

17. I. Schmitt, M. Linden, H. Khazneh, B. O. Evert, P. Breuer, T. Klockgether, U. Wuellner, Inactivation of the mouse Atxn3 (ataxin-3) gene increases protein ubiquitination. Biochem Biophys Res Commun 362, 734–739 (2007).

18. K. H. Schuster, D. M. DiFranco, A. F. Putka, J. P. Mato, S. I. Jarrah, N. R. Stec, V. O. Sundararajan, H. S. McLoughlin, Disease-associated oligodendrocyte signatures are spatiotemporally dysregulated in spinocerebellar ataxia type 3. Front Neurosci 17, 1118429 (2023).

19. K. H. Schuster, A. F. Putka, H. S. McLoughlin, Pathogenetic Mechanisms Underlying Spinocerebellar Ataxia Type 3 Are Altered in Primary Oligodendrocyte Culture. Cells 11, 2615 (2022).

20. J. Faber, T. Schaprian, K. Berkan, K. Reetz, M. C. França, T. J. R. de Rezende, J. Hong, W. Liao, B. van de Warrenburg, J. van Gaalen, A. Durr, F. Mochel, P. Giunti, H. Garcia-Moreno, L. Schoels, H. Hengel, M. Synofzik, B. Bender, G. Oz, J. Joers, J. J. de Vries, J.-S. Kang, D. Timmann-Braun, H. Jacobi, J. Infante, R. Joules, S. Romanzetti, J. Diedrichsen, M. Schmid, R. Wolz, T. Klockgether, Regional Brain and Spinal Cord Volume Loss in Spinocerebellar Ataxia Type 3. Mov Disord 36, 2273–2281 (2021).

21. W. O. Arruda, A. T. Meira, S. E. Ono, A. de Carvalho Neto, L. E. G. G. Betting, S. Raskin, C. H. F. Camargo, H. A. G. Teive, Volumetric MRI Changes in Spinocerebellar Ataxia (SCA3 and SCA10) Patients. Cerebellum 19, 536–543 (2020).

22. J. Chandrasekaran, E. Petit, Y.-W. Park, S. Tezenas du Montcel, J. M. Joers, D. K. Deelchand, M. Považan, G. Banan, R. Valabregue, P. Ehses, J. Faber, P. Coupé, C. U. Onyike, P. B. Barker, J. D. Schmahmann, E.-M. Ratai, S. H. Subramony, T. H. Mareci, K. O. Bushara, H. Paulson, A. Durr, T. Klockgether, T. Ashizawa, C. Lenglet, G. Öz, Clinically meaningful MR endpoints sensitive to preataxic SCA1 and SCA3. Ann Neurol 93, 686–701 (2023).

23. J.-S. Kang, J. C. Klein, S. Baudrexel, R. Deichmann, D. Nolte, R. Hilker, White matter damage is related to ataxia severity in SCA3. J Neurol 261, 291–299 (2014).

24. X. Wu, X. Liao, Y. Zhan, C. Cheng, W. Shen, M. Huang, Z. Zhou, Z. Wang, Z. Qiu, W. Xing, W. Liao, B. Tang, L. Shen, Microstructural Alterations in Asymptomatic and Symptomatic Patients with Spinocerebellar Ataxia Type 3: A Tract-Based Spatial Statistics Study. Front Neurol 8, 714 (2017).

25. M. do C. Costa, M. Radzwion, H. S. McLoughlin, N. S. Ashraf, S. Fischer, V. G. Shakkottai, P. Maciel, H. L. Paulson, G. Öz, In Vivo Molecular Signatures of Cerebellar Pathology in Spinocerebellar Ataxia Type 3. Movement Disorders 35, 1774–1786 (2020).

26. K. H. Schuster, A. J. Zalon, D. M. DiFranco, A. F. Putka, N. R. Stec, S. I. Jarrah, A. Naeem, Z. Haque, H. Zhang, Y. Guan, H. S. McLoughlin, ASOs are an effective treatment for disease-associated oligodendrocyte signatures in premanifest and symptomatic SCA3 mice. Molecular Therapy 32, 1359–1372 (2024).

27. N. Baumann, D. Pham-Dinh, Biology of oligodendrocyte and myelin in the mammalian central nervous system. Physiol Rev 81, 871–927 (2001).

28. S. E. Pfeiffer, A. E. Warrington, R. Bansal, The oligodendrocyte and its many cellular processes. Trends Cell Biol 3, 191–197 (1993).

29. W. Lin, B. Popko, Endoplasmic reticulum stress in disorders of myelinating cells. Nat Neurosci 12, 379–385 (2009).

30. M. C. Truttmann, V. E. Cruz, X. Guo, C. Engert, T. U. Schwartz, H. L. Ploegh, The Caenorhabditis elegans Protein FIC-1 Is an AMPylase That Covalently Modifies Heat-Shock 70 Family Proteins, Translation Elongation Factors and Histones. PLoS Genet 12, e1006023 (2016).

31. M. C. Truttmann, H. L. Ploegh, rAMPing Up Stress Signaling: Protein AMPylation in Metazoans. Trends Cell Biol. 27, 608–620 (2017).

32. S. Preissler, C. Rato, R. Chen, R. Antrobus, S. Ding, I. M. Fearnley, D. Ron, AMPylation matches BiP activity to client protein load in the endoplasmic reticulum. Elife 4, e12621 (2015).

33. H. Ham, A. R. Woolery, C. Tracy, D. Stenesen, H. Krämer, K. Orth, Unfolded protein response-regulated Drosophila Fic (dFic) protein reversibly AMPylates BiP chaperone during endoplasmic reticulum homeostasis. J. Biol. Chem. 289, 36059–36069 (2014).

34. A. Sanyal, A. J. Chen, E. S. Nakayasu, C. S. Lazar, E. A. Zbornik, C. A. Worby, A. Koller, S. Mattoo, A novel link between Fic (filamentation induced by cAMP)-mediated adenylylation/AMPylation and the unfolded protein response. J. Biol. Chem. 290, 8482– 8499 (2015).

35. M. C. Truttmann, X. Zheng, L. Hanke, J. R. Damon, M. Grootveld, J. Krakowiak, D. Pincus, H. L. Ploegh, Unrestrained AMPylation targets cytosolic chaperones and activates the heat shock response. Proc. Natl. Acad. Sci. U.S.A. 114, E152–E160 (2017).

36. L. A. Perera, C. Rato, Y. Yan, L. Neidhardt, S. H. McLaughlin, R. J. Read, S. Preissler, D. Ron, An oligomeric state-dependent switch in the ER enzyme FICD regulates AMPylation and deAMPylation of BiP. The EMBO Journal 38, e102177 (2019).

37. L. A. Perera, S. Preissler, N. R. Zaccai, S. Prévost, J. M. Devos, M. Haertlein, D. Ron, Structures of a deAMPylation complex rationalise the switch between antagonistic catalytic activities of FICD. Nat Commun 12, 5004 (2021).

38. S. Preissler, C. Rato, L. Perera, V. Saudek, D. Ron, FICD acts bifunctionally to AMPylate and de-AMPylate the endoplasmic reticulum chaperone BiP. Nat Struct Mol Biol 24, 23–29 (2017).

39. L. Wieteska, S. Shahidi, A. Zhuravleva, Allosteric fine-tuning of the conformational equilibrium poises the chaperone BiP for post-translational regulation. Elife 6, e29430 (2017).

40. L. A. Perera, A. T. Hattersley, H. P. Harding, M. N. Wakeling, S. E. Flanagan, I. Mohsina, J. Raza, A. Gardham, D. Ron, E. De Franco, Infancy-onset diabetes caused by de-regulated AMPylation of the human endoplasmic reticulum chaperone BiP. EMBO Mol Med 15, e16491 (2023).

41. S. S. Hassan, S. A. Musa, E. De Franco, R. Myers, R. Van Heugten, O. O. Babiker, A. A. Ibrahim, G. F. MohamadSalih, A. Ahmed, J. A. Shatta, O. A. Al-Hassan, K. A. Patel, M. A. Abdullah, Characterization of monogenic diabetes among Sudanese children: a multi-center experience from a population with high consanguinity. J Pediatr Endocrinol Metab 39, 65–75.

42. A. P. Rebelo, A. Ruiz, M. F. Dohrn, M. Wayand, A. Farooq, M. C. Danzi, D. Beijer, B. Aaron, J. Vandrovcova, H. Houlden, L. Matalonga, L. Abreu, G. Rouleau, M. A. Estiar, L. Van de Vondel, Z. Gan-Or, J. Baets, R. Schüle, S. Zuchner, BiP inactivation due to loss of the deAMPylation function of FICD causes a motor neuron disease. Genet Med 24, 2487– 2500 (2022).

43. M. Vinci, D. Greco, M. G. Figura, S. Treccarichi, A. Musumeci, V. Greco, R. Pettinato, A. Gloria, C. Papa, S. Saccone, C. Federico, F. Calì, Exploring the Role of FICD, a New Potential Gene Involved in Borderline Intellectual Functioning, Psychological and Metabolic Disorders. Genes 15, 1655 (2024).

44. N. McCaul, C. M. Porter, A. Becker, C.-H. A. Tang, C. Wijne, B. Chatterjee, D. Bousbaine, A. Bilate, C.-C. A. Hu, H. Ploegh, M. C. Truttmann, Deletion of mFICD AMPylase alters cytokine secretion and affects visual short-term learning in vivo. J Biol Chem 297, 100991 (2021).

45. A. K. Casey, H. F. Gray, S. Chimalapati, G. Hernandez, A. T. Moehlman, N. Stewart, H. A. Fields, B. Gulen, K. A. Servage, K. Stefanius, A. Blevins, B. M. Evers, H. Krämer, K. Orth, Fic-mediated AMPylation tempers the unfolded protein response during physiological stress. Proc Natl Acad Sci U S A 119, e2208317119 (2022).

46. A. K. Casey, N. M. Stewart, N. Zaidi, H. F. Gray, H. A. Fields, M. Sakurai, C. A. Pinzon-Arteaga, B. M. Evers, J. Wu, K. Orth, Pre-clinical model of dysregulated FicD AMPylation causes diabetes by disrupting pancreatic endocrine homeostasis. Mol Metab 95, 102120 (2025).

47. S. M. Lacy, R. J. Taubitz, N. D. Urban, S. N. Turowski, E. N. Primack, E. D. Smith, A. S. Helms, D. E. Michele, M. C. Truttmann, FICD (FIC Domain Protein Adenylyl Transferase) Deficiency Protects Mice From Hypertrophy-Induced Heart Failure and Promotes BiP (Binding Immunoglobulin Protein) -Mediated Activation of the Unfolded Protein Response and Endoplasmic Reticulum-Selective Autophagy in Cardiomyocytes. J Am Heart Assoc 14, e040192 (2025).

48. M. C. Truttmann, D. Pincus, H. L. Ploegh, Chaperone AMPylation modulates aggregation and toxicity of neurodegenerative disease-associated polypeptides. Proc. Natl. Acad. Sci. U.S.A. 115, E5008–E5017 (2018).

49. A. Sanyal, S. Dutta, A. Camara, A. Chandran, A. Koller, B. G. Watson, R. Sengupta, D. Ysselstein, P. Montenegro, J. Cannon, J.-C. Rochet, S. Mattoo, Alpha-Synuclein Is a Target of Fic-Mediated Adenylylation/AMPylation: Possible Implications for Parkinson’s Disease. J Mol Biol 431, 2266–2282 (2019).

50. A. Koller, L. Hoffmann, A. Bluhm, A. Schweigert, Y. Schneider, M. Andert, T. Becker, F. Zunke, T. G. Beach, G. E. Serrano, S. Roßner, J. Winkler, P. Kielkowski, W. Xiang, Aberrant FICD-mediated AMPylation drives α-Synuclein pathology and overall protein dyshomeostasis in dopaminergic neurons in Parkinson’s disease. bioRxiv, 2026.03.30.715195 (2026).

51. K. M. Van Pelt, M. C. Truttmann, Loss of FIC-1-mediated AMPylation activates the UPRER and upregulates cytosolic HSP70 chaperones to suppress polyglutamine toxicity. PLoS Genet 21, e1011723 (2025).

52. Y. Hussien, J. R. Podojil, A. P. Robinson, A. S. Lee, S. D. Miller, B. Popko, ER Chaperone BiP/GRP78 Is Required for Myelinating Cell Survival and Provides Protection during Experimental Autoimmune Encephalomyelitis. J Neurosci 35, 15921–15933 (2015).

53. W.-C. M. Lee, M. Yoshihara, J. T. Littleton, Cytoplasmic aggregates trap polyglutamine-containing proteins and block axonal transport in a Drosophila model of Huntington’s disease. Proceedings of the National Academy of Sciences 101, 3224–3229 (2004).

54. B. Eftekharzadeh, A. Piai, G. Chiesa, D. Mungianu, J. García, R. Pierattelli, I. C. Felli, X. Salvatella, Sequence Context Influences the Structure and Aggregation Behavior of a PolyQ Tract. Biophysical Journal 110, 2361–2366 (2016).

55. C. K. Cemal, C. J. Carroll, L. Lawrence, M. B. Lowrie, P. Ruddle, S. Al-Mahdawi, R. H. M. King, M. A. Pook, C. Huxley, S. Chamberlain, YAC transgenic mice carrying pathological alleles of the MJD1 locus exhibit a mild and slowly progressive cerebellar deficit. Hum Mol Genet 11, 1075–1094 (2002).

56. M. do C. Costa, K. Luna-Cancalon, S. Fischer, N. S. Ashraf, M. Ouyang, R. M. Dharia, L. Martin-Fishman, Y. Yang, V. G. Shakkottai, B. L. Davidson, E. Rodríguez-Lebrón, H. L. Paulson, Toward RNAi therapy for the polyglutamine disease Machado-Joseph disease. Mol Ther 21, 1898–1908 (2013).

57. V. G. Shakkottai, M. do Carmo Costa, J. M. Dell’Orco, A. Sankaranarayanan, H. Wulff, H. L. Paulson, Early changes in cerebellar physiology accompany motor dysfunction in the polyglutamine disease spinocerebellar ataxia type 3. J Neurosci 31, 13002–13014 (2011).

58. D. Höpfner, J. Fauser, M. S. Kaspers, C. Pett, C. Hedberg, A. Itzen, Monoclonal Anti-AMP Antibodies Are Sensitive and Valuable Tools for Detecting Patterns of AMPylation. iScience 23, 101800 (2020).

59. R. M. Nowier, A. Friedman, A. Brown, P. Jung, The role of neurofilament transport in the radial growth of myelinated axons. Mol Biol Cell 34, ar58 (2023).

60. S. Aggarwal, L. Yurlova, M. Simons, Central nervous system myelin: structure, synthesis and assembly. Trends Cell Biol 21, 585–593 (2011).

61. J. M. Bin, S. N. Harris, T. E. Kennedy, The oligodendrocyte-specific antibody ‘CC1’ binds Quaking 7. Journal of Neurochemistry 139, 181–186 (2016).

62. G. Saher, B. Brügger, C. Lappe-Siefke, W. Möbius, R. Tozawa, M. C. Wehr, F. Wieland, S. Ishibashi, K.-A. Nave, High cholesterol level is essential for myelin membrane growth. Nat Neurosci 8, 468–475 (2005).

63. M. Orth, S. Bellosta, Cholesterol: Its Regulation and Role in Central Nervous System Disorders. Cholesterol 2012, 292598 (2012).

64. B. B. Madison, Srebp2: A master regulator of sterol and fatty acid synthesis1. J Lipid Res 57, 333–335 (2016).

65. N.-Q. Wang, P.-X. Sun, Q.-Q. Shen, M.-Y. Deng, Cholesterol Metabolism in CNS Diseases: The Potential of SREBP2 and LXR as Therapeutic Targets. Mol Neurobiol 62, 6283–6307 (2025).

66. K. L. Cook, D. R. Soto-Pantoja, P. A. G. Clarke, M. I. Cruz, A. Zwart, A. Wärri, L. Hilakivi-Clarke, D. D. Roberts, R. Clarke, Endoplasmic reticulum stress protein GRP78 modulates lipid metabolism to control drug sensitivity and anti-tumor immunity in breast cancer. Cancer Res 76, 5657–5670 (2016).

67. B. Huang, W. Wei, G. Wang, M. A. Gaertig, Y. Feng, W. Wang, X.-J. Li, S. Li, Mutant huntingtin downregulates myelin regulatory factor-mediated myelin gene expression and affects mature oligodendrocytes. Neuron 85, 1212–1226 (2015).

68. C. Ferrari Bardile, M. Garcia-Miralles, N. S. Caron, N. A. Rayan, S. R. Langley, N. Harmston, A. M. Rondelli, R. T. Y. Teo, S. Waltl, L. M. Anderson, H.-G. Bae, S. Jung, A. Williams, S. Prabhakar, E. Petretto, M. R. Hayden, M. A. Pouladi, Intrinsic mutant HTT-mediated defects in oligodendroglia cause myelination deficits and behavioral abnormalities in Huntington disease. Proc Natl Acad Sci U S A 116, 9622–9627 (2019).

69. C. Lee, R. M. Grijalva, L. Tejwani, E. Bae, A. Chase, H. Ro, H. Kim, V. Olmos, J. P. Orengo, J. Lim, Oligodendrocyte dysfunction contributes to motor deficits and Purkinje cell axonopathy in spinocerebellar ataxia type 1. J Clin Invest 136 (2026).

70. C. Lukas, L. Schöls, B. Bellenberg, U. Rüb, H. Przuntek, G. Schmid, O. Köster, B. Suchan, Dissociation of grey and white matter reduction in spinocerebellar ataxia type 3 and 6: a voxel-based morphometry study. Neurosci Lett 408, 230–235 (2006).

71. A. D’Abreu, M. C. França, H. L. Paulson, I. Lopes-Cendes, Caring for Machado–Joseph disease: Current understanding and how to help patients. Parkinsonism & Related Disorders 16, 2–7 (2010).

72. P. Kielkowski, I. Y. Buchsbaum, V. C. Kirsch, N. C. Bach, M. Drukker, S. Cappello, S. A. Sieber, FICD activity and AMPylation remodelling modulate human neurogenesis. Nat Commun 11, 517 (2020).

73. L. Hoffmann, E.-M. Eckl, M. Bérouti, M. Pries, A. Koller, C. Guhl, U. A. Hellmich, V. Hornung, W. Xiang, L. T. Jae, P. Kielkowski, AMPylation Regulates 5’-3’ Exonuclease PLD3 Processing. Mol Cell Proteomics 24, 101051 (2025).

74. A. K. Casey, N. M. Stewart, N. Zaidi, H. F. Gray, A. Cox, H. A. Fields, K. Orth, FicD regulates adaptation to the unfolded protein response in the murine liver. Biochimie 225, 114–124 (2024).

75. E. Hernández-Carralero, G. Quinet, R. Freire, ATXN3: a multifunctional protein involved in the polyglutamine disease spinocerebellar ataxia type 3. Expert Rev Mol Med 26, e19 (2024).

76. X. Zhong, R. N. Pittman, Ataxin-3 binds VCP/p97 and regulates retrotranslocation of ERAD substrates. Human Molecular Genetics 15, 2409–2420 (2006).

77. M. Boyer-Guittaut, L. Poillet, Q. Liang, E. Bôle-Richard, X. Ouyang, G. A. Benavides, F.-Z. Chakrama, A. Fraichard, V. M. Darley-Usmar, G. Despouy, M. Jouvenot, R. Delage-Mourroux, J. Zhang, The role of GABARAPL1/GEC1 in autophagic flux and mitochondrial quality control in MDA-MB-436 breast cancer cells. Autophagy 10, 986– 1003 (2014).

78. Z. Sha, H. M. Schnell, K. Ruoff, A. Goldberg, Rapid induction of p62 and GABARAPL1 upon proteasome inhibition promotes survival before autophagy activation. J Cell Biol 217, 1757–1776 (2018).

79. E. D. Evalt, S. Govindaraj, M. T. Jones, N. Ozsoy, H. Chen, A. E. Russell, Endoplasmic reticulum stress alters myelin associated protein expression and extracellular vesicle composition in human oligodendrocytes. Front Mol Biosci 11, 1432945 (2024).

80. B. L. L. Clayton, B. Popko, Endoplasmic reticulum stress and the unfolded protein response in disorders of myelinating glia. Brain Res 1648, 594–602 (2016).

81. Y. Gao, L. P. Slomnicki, E. Kilanczyk, M. D. Forston, M. Pietrzak, E. C. Rouchka, R. M. Howard, S. R. Whittemore, M. Hetman, Reduced Expression of Oligodendrocyte Linage-Enriched Transcripts During the Endoplasmic Reticulum Stress/Integrated Stress Response. ASN Neuro 16, 2371162 (2024).

82. S. S. Ohri, M. Hetman, S. R. Whittemore, Restoring endoplasmic reticulum homeostasis improves functional recovery after spinal cord injury. Neurobiology of Disease 58, 29–37 (2013).

83. M. Valenza, D. Rigamonti, D. Goffredo, C. Zuccato, S. Fenu, L. Jamot, A. Strand, A. Tarditi, B. Woodman, M. Racchi, C. Mariotti, S. Di Donato, A. Corsini, G. Bates, R. Pruss, J. M. Olson, S. Sipione, M. Tartari, E. Cattaneo, Dysfunction of the Cholesterol Biosynthetic Pathway in Huntington’s Disease. J Neurosci 25, 9932–9939 (2005).

84. A. F. Putka, V. Mohanty, S. M. Cologna, H. S. McLoughlin, Cerebellar Lipid Dysregulation in SCA3: A Comparative Study in Patients and Mice. Neurobiol Dis 206, 106827 (2025).

85. C. Nóbrega, L. Mendonça, A. Marcelo, A. Lamazière, S. Tomé, G. Despres, C. A. Matos, F. Mechmet, D. Langui, W. den Dunnen, L. P. de Almeida, N. Cartier, S. Alves, Restoring brain cholesterol turnover improves autophagy and has therapeutic potential in mouse models of spinocerebellar ataxia. Acta Neuropathol 138, 837–858 (2019).

86. B. K. Chatterjee, M. Alam, A. Chakravorty, S. M. Lacy, W. Giblin, J. Rech, C. L. I. Brooks, P. Arvan, M. C. Truttmann, Small-Molecule FICD Inhibitors Suppress Endogenous and Pathologic FICD-Mediated Protein AMPylation. ACS Chem. Biol. 20, 880–895 (2025).

87. E. Sjöstedt, W. Zhong, L. Fagerberg, M. Karlsson, N. Mitsios, C. Adori, P. Oksvold, F. Edfors, A. Limiszewska, F. Hikmet, J. Huang, Y. Du, L. Lin, Z. Dong, L. Yang, X. Liu, H. Jiang, X. Xu, J. Wang, H. Yang, L. Bolund, A. Mardinoglu, C. Zhang, K. von Feilitzen, C. Lindskog, F. Pontén, Y. Luo, T. Hökfelt, M. Uhlén, J. Mulder, An atlas of the protein-coding genes in the human, pig, and mouse brain. Science 367, eaay5947 (2020).

88. K. J. E. Matson, D. E. Russ, C. Kathe, I. Hua, D. Maric, Y. Ding, J. Krynitsky, R. Pursley, A. Sathyamurthy, J. W. Squair, B. P. Levi, G. Courtine, A. J. Levine, Single cell atlas of spinal cord injury in mice reveals a pro-regenerative signature in spinocerebellar neurons. Nat Commun 13, 5628 (2022).

89. L. Khandker, M. A. Jeffries, Y.-J. Chang, M. L. Mather, A. V. Evangelou, J. N. Bourne, A. K. Tafreshi, I. M. Ornelas, O. Bozdagi-Gunal, W. B. Macklin, T. L. Wood, Cholesterol biosynthesis defines oligodendrocyte precursor heterogeneity between brain and spinal cord. Cell Rep 38, 110423 (2022).

90. L. R. Moore, G. Rajpal, I. T. Dillingham, M. Qutob, K. G. Blumenstein, D. Gattis, G. Hung, H. B. Kordasiewicz, H. L. Paulson, H. S. McLoughlin, Evaluation of Antisense Oligonucleotides Targeting ATXN3 in SCA3 Mouse Models. Mol Ther Nucleic Acids 7, 200–210 (2017).

91. J. Schindelin, I. Arganda-Carreras, E. Frise, V. Kaynig, M. Longair, T. Pietzsch, S. Preibisch, C. Rueden, S. Saalfeld, B. Schmid, J.-Y. Tinevez, D. J. White, V. Hartenstein, K. Eliceiri, P. Tomancak, A. Cardona, Fiji: an open-source platform for biological-image analysis. Nat Methods 9, 676–682 (2012).

92. A. T. Kong, F. V. Leprevost, D. M. Avtonomov, D. Mellacheruvu, A. I. Nesvizhskii, MSFragger: ultrafast and comprehensive peptide identification in mass spectrometry-based proteomics. Nat Methods 14, 513–520 (2017).

93. F. Yu, G. C. Teo, A. T. Kong, K. Fröhlich, G. X. Li, V. Demichev, A. I. Nesvizhskii, Analysis of DIA proteomics data using MSFragger-DIA and FragPipe computational platform. Nat Commun 14, 4154 (2023).

94. K. L. Yang, F. Yu, G. C. Teo, K. Li, V. Demichev, M. Ralser, A. I. Nesvizhskii, MSBooster: improving peptide identification rates using deep learning-based features. Nat Commun 14, 4539 (2023).

95. L. Käll, J. D. Canterbury, J. Weston, W. S. Noble, M. J. MacCoss, Semi-supervised learning for peptide identification from shotgun proteomics datasets. Nat Methods 4, 923– 925 (2007).

96. A. I. Nesvizhskii, A. Keller, E. Kolker, R. Aebersold, A statistical model for identifying proteins by tandem mass spectrometry. Anal Chem 75, 4646–4658 (2003).

97. F. da Veiga Leprevost, S. E. Haynes, D. M. Avtonomov, H.-Y. Chang, A. K. Shanmugam, D. Mellacheruvu, A. T. Kong, A. I. Nesvizhskii, Philosopher: a versatile toolkit for shotgun proteomics data analysis. Nat Methods 17, 869–870 (2020).

98. V. Demichev, C. B. Messner, S. I. Vernardis, K. S. Lilley, M. Ralser, DIA-NN: neural networks and interference correction enable deep proteome coverage in high throughput. Nat Methods 17, 41–44 (2020).

99. Y. Hsiao, H. Zhang, G. X. Li, Y. Deng, F. Yu, H. Valipour Kahrood, J. R. Steele, R. B. Schittenhelm, A. I. Nesvizhskii, Analysis and Visualization of Quantitative Proteomics Data Using FragPipe-Analyst. J Proteome Res 23, 4303–4315 (2024).

100. Y. Jiao, Z. Sun, T. Lee, F. R. Fusco, T. D. Kimble, C. A. Meade, S. Cuthbertson, A. Reiner, A simple and sensitive antigen retrieval method for free-floating and slide-mounted tissue sections. J Neurosci Methods 93, 149–162 (1999).

101. C. Stringer, M. Pachitariu, Cellpose3: one-click image restoration for improved cellular segmentation. Nat Methods 22, 592–599 (2025).

102. D. R. Stirling, M. J. Swain-Bowden, A. M. Lucas, A. E. Carpenter, B. A. Cimini, A. Goodman, CellProfiler 4: improvements in speed, utility and usability. BMC Bioinformatics 22, 433 (2021).

103. E. Weisbart, C. Tromans-Coia, B. Diaz-Rohrer, D. R. Stirling, F. Garcia-Fossa, R. A. Senft, M. C. Hiner, M. B. de Jesus, K. W. Eliceiri, B. A. Cimini, CellProfiler plugins - An easy image analysis platform integration for containers and Python tools. J Microsc 296, 227–234 (2024).

104. F. Xie, J. Wang, B. Zhang, RefFinder: a web-based tool for comprehensively analyzing and identifying reference genes. Funct Integr Genomics 23, 125 (2023).

105. Y. Chen, L. Chen, A. T. L. Lun, P. L. Baldoni, G. K. Smyth, edgeR v4: powerful differential analysis of sequencing data with expanded functionality and improved support for small counts and larger datasets. Nucleic Acids Res 53, gkaf018 (2025).

106. T. Wu, E. Hu, S. Xu, M. Chen, P. Guo, Z. Dai, T. Feng, L. Zhou, W. Tang, L. Zhan, X. Fu, S. Liu, X. Bo, G. Yu, clusterProfiler 4.0: A universal enrichment tool for interpreting omics data. Innovation 2 (2021).

107. H. L. Paulson, S. S. Das, P. B. Crino, M. K. Perez, S. C. Patel, D. Gotsdiner, K. H. Fischbeck, R. N. Pittman, Machado-Joseph disease gene product is a cytoplasmic protein widely expressed in brain. Annals of Neurology 41, 453–462 (1997).

108. S. Smith, E. R. Swan, K. L. Furber, Establishing validated RT-qPCR workflow for the analysis of oligodendrocyte gene expression in the developing murine brain. Biochem Cell Biol 102, 492–505 (2024).

109. W. Y. Ho, J.-C. Chang, K. Lim, A. Cazenave-Gassiot, A. T. Nguyen, J. C. Foo, S. Muralidharan, A. Viera-Ortiz, S. J. M. Ong, J. H. Hor, I. Agrawal, S. Hoon, O. A. Arogundade, M. J. Rodriguez, S. M. Lim, S. H. Kim, J. Ravits, S.-Y. Ng, M. R. Wenk, E. B. Lee, G. Tucker-Kellogg, S.-C. Ling, TDP-43 mediates SREBF2-regulated gene expression required for oligodendrocyte myelination. J Cell Biol 220, e201910213 (2021).

